# PrkC modulates MreB filament density and cellular growth rate by monitoring cell wall precursors

**DOI:** 10.1101/2020.08.28.272336

**Authors:** Yingjie Sun, Ethan Garner

## Abstract

How bacteria link their rate of growth to the external nutrient conditions is not known. To explore how *Bacillus subtilis* modulates the rate cells expand their encapsulating cell wall, we compared the single-cell growth rate to the density of moving MreB filaments under different conditions. MreB filament density scales with the growth rate, and is modulated by the *mur* genes that create the cell wall precursor lipid II. Lipid II is sensed by the serine/threonine kinase PrkC, which, among other proteins, phosphorylates RodZ. Phosphorylated RodZ then increases MreB filament density, increasing growth. Strikingly, increasing the activity of this pathway results in cells elongating far faster than wild type in nutrient-poor media, indicating slow-growing bacteria contain spare growth capacity. Overall, this work reveals that PrkC functions as a cellular rheostat, tuning the activities of cellular processes in response to lipid II, allowing cells to grow robustly across a broad range of nutrient conditions.

**One-sentence summary:** The serine/threonine kinase PrkC modulates both MreB filament density and cellular growth rate by sensing lipid II in *Bacillus subtilis*.

## Main Text

Since the seminal work of Monod, bacteria have been held to grow as fast as the most limiting nutrient within the media allows (Monod 1949). To accomplish this, bacteria must balance the rates they synthesize their essential components: their RNA/protein ratio (Kjeldgaard, Maaloe, and Schaechter 1958; Schaechter, Maaloe, and Kjeldgaard 1958), DNA replication frequency, and the rates they synthesize membranes and build external protective structures. For bacteria, a critical essential component is their peptidoglycan (PG) cell wall, a covalently crosslinked meshwork that surrounds cells conferring both shape and protection from lysis. In order to grow and divide, bacteria cells must add new material into this protective structure.

The gram-positive *Bacillus subtilis* inserts new PG into the wall via the action of two different PG synthetic systems: the Class A PBPs and the Rod complex (Cho et al. 2016; Meeske et al. 2016). The Rod complex contains proteins required for rod shape: the transglycosylase/transpeptidase pair RodA/PBP2a, MreC, MreD, RodZ, and MreB (Fig. 1A). MreB polymerizes into short, curved membrane-associated filaments that orient to point around the rod circumference (Hussain et al. 2018). MreB orientation constrains the Rod complex to move around the rod circumference as it synthesizes new PG (Garner et al. 2011; Dominguez-Escobar et al. 2011), resulting in a radial arrangement of glycans that reinforce the cells rod shape against internal turgor (Dion et al. 2019). It is not known how bacteria regulate the rates PG synthetic enzymes build new cell wall, much less how they scale their activity to match the synthesis of other biomolecules. To explore these questions, we sought to understand how PG synthetic activity scaled with the growth rate in *B. subtilis*, subsequently exploring the mechanisms by which this regulation occurs.

**Fig. 1.**
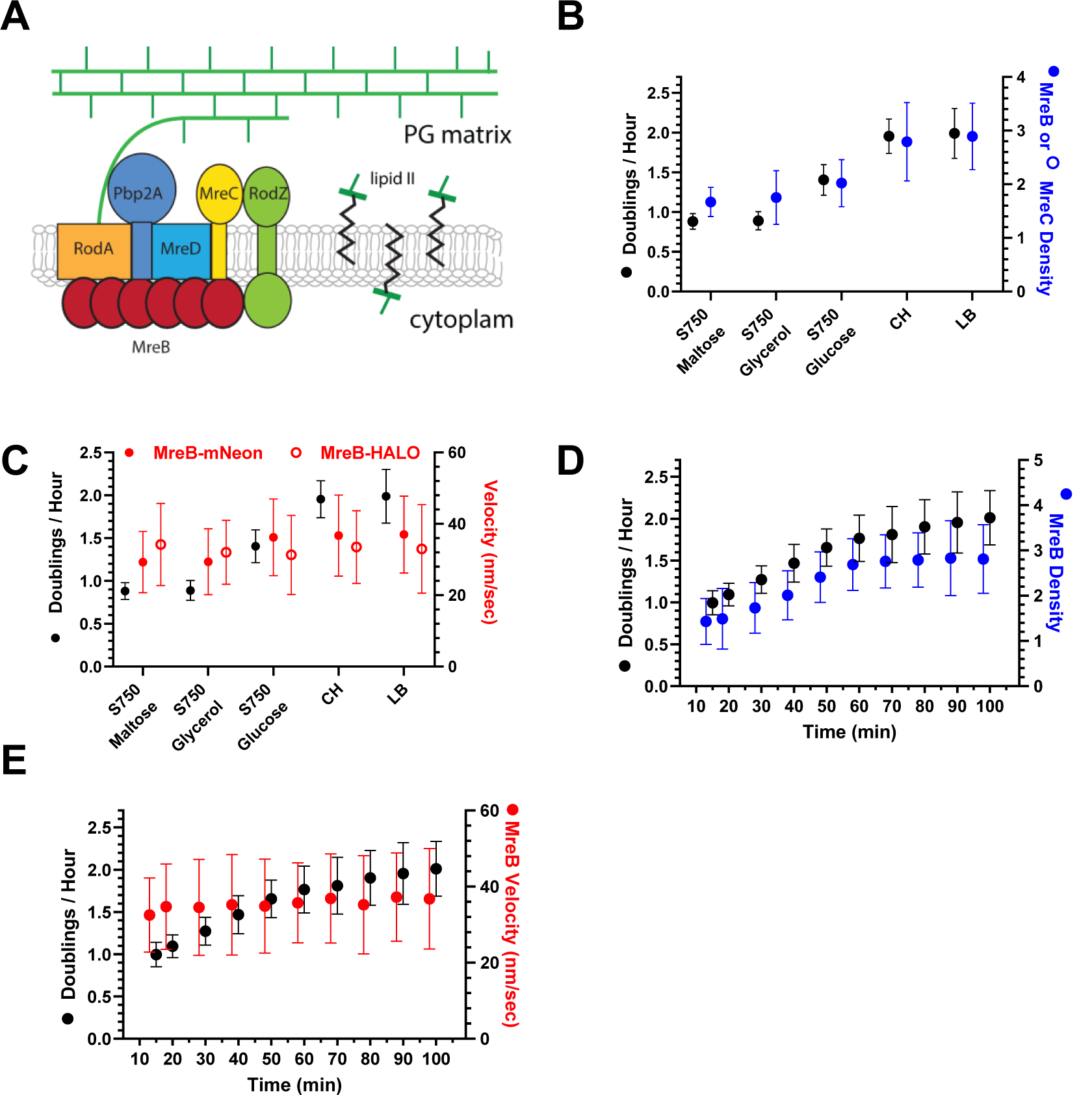
MreB filament density - and not velocity - increases with growth rate. **(A) Schematic of Rod complex** **(B) Directional MreB and MreC density increases with the growth rate -** Cells expressing MreB-mNeonGreen (bYS09) or MreC-mNeonGreen (bYS170) were grown in the indicated media under agar pads, and their growth rate (left axis), and directional MreB and MreC density (right axis) determined by single-cell analysis. See also Fig. S1A and Movies S1 and S3. **(C) MreB velocity remains the same in different media -** Cells expressing MreB-mNeonGreen (bYS09) were grown in different media, and MreB velocity determined by particle tracking and MSD analysis. See also S1B and Movie S1-2. **(D to E) MreB filament density, but not velocity increases during nutrient upshifts** - Cells were grown in S750 maltose for 6 hours in liquid, washed in CH, then placed under a pad made of CH media immediately before imaging with TIRF-SIM. Plotted is the growth rate and either the density (**D**) or velocity (**E**) of directionally moving MreB filaments. See also Movies S4 and S5.

Given the directional motions of the Rod Complex are driven by PG synthesis (Garner et al. 2011; Dominguez-Escobar et al. 2011), their activity can be measured by quantifying the number and velocity of directionally moving MreB filaments. As filaments might be too dense to resolve using diffraction-limited microscopy, we used a method that counts the number of directionally moving particles using temporal correlations between adjacent pixels. This approach is more accurate than particle tracking, giving measures equivalent to tracking filaments imaged with TIRF-structured illumination microscopy (TIRF-SIM) (Dion et al. 2019).

We started by measuring the density and velocity of MreB filaments in cells growing under agar pads composed of different media, along with their single-cell growth rate (Fig. 1B-C). In contrast to previous reports that tracked MreB filaments imaged with TIRF (Billaudeau et al. 2017), we find that filament density increases with growth rate (Fig. 1B-C). We verified this observation using TIRF-SIM (Movie S1); cells growing in rich media contained numerous filaments, many of which were small and sub-diffraction in size, and filament density decreased as cells were grown in progressively less rich media. MreB filament velocity remained mostly constant across growth conditions (Fig. 1C), a finding further verified by tracking single MreB-HaloTag molecules (Fig. 1C, Movie S2). Similar results were obtained in cells expressing msfGFP fusions to both MreB and Mbl (Fig. S1A,B). We believe the differences between this and the Billaudeau study likely arises due to the different fluorescent fusions to MreB (Supplemental Text 1). Another Rod complex component showed the same trend: the number and velocity of MreC-mNeonGreen molecules gave nearly identical measures of directionally moving particles as MreB under different growth conditions, increasing in number (Fig. 1B) but invariant in velocity (Fig. S1C). Interestingly, during slow growth most MreC-mNeonGreen molecules were diffusive, and as growth rate increased, more MreC molecules moved directionally (Movie S3). Likewise, when cells were shifted from slow growth media to fast growth media, filament number increased while velocity remained constant (Fig. 1D-E, Movies S4 and S5). Thus, *B. subtilis* increases the number of directionally moving Rod complexes to increase its growth rate.

Next, we explored how cells control the number of active Rod complexes. This regulation is unlikely to occur via changes in expression, as proteomics suggest most Rod Complex components remain relatively constant across different growth media (Fig. S2A). Alternatively, past work suggests a role for lipid II, the immediate precursor for PG synthesis: not only does the relative abundance of Mur proteins scale with growth rate (Fig. S2B) (Dion et al. 2019), perturbations that deplete cells of lipid II cause MreB filaments to depolymerize off of the membrane, inactivating the associated synthetic enzymes (Schirner et al. 2015). To test if lipid II production affected MreB filament density, we titrated the expression of the first enzyme in the Mur pathway, *murAA*. In nutrient-rich media, low *murAA* inductions decreased the growth rate, and also MreB and MreC density (Fig. 2A, B). As induction increased, so did growth rate and MreB and MreC density, with high inductions reaching wild type (WT) values. As before, MreB and MreC velocity remained unchanged (Fig. 2C, Movie S6, and S2C).

**Fig. 2.**
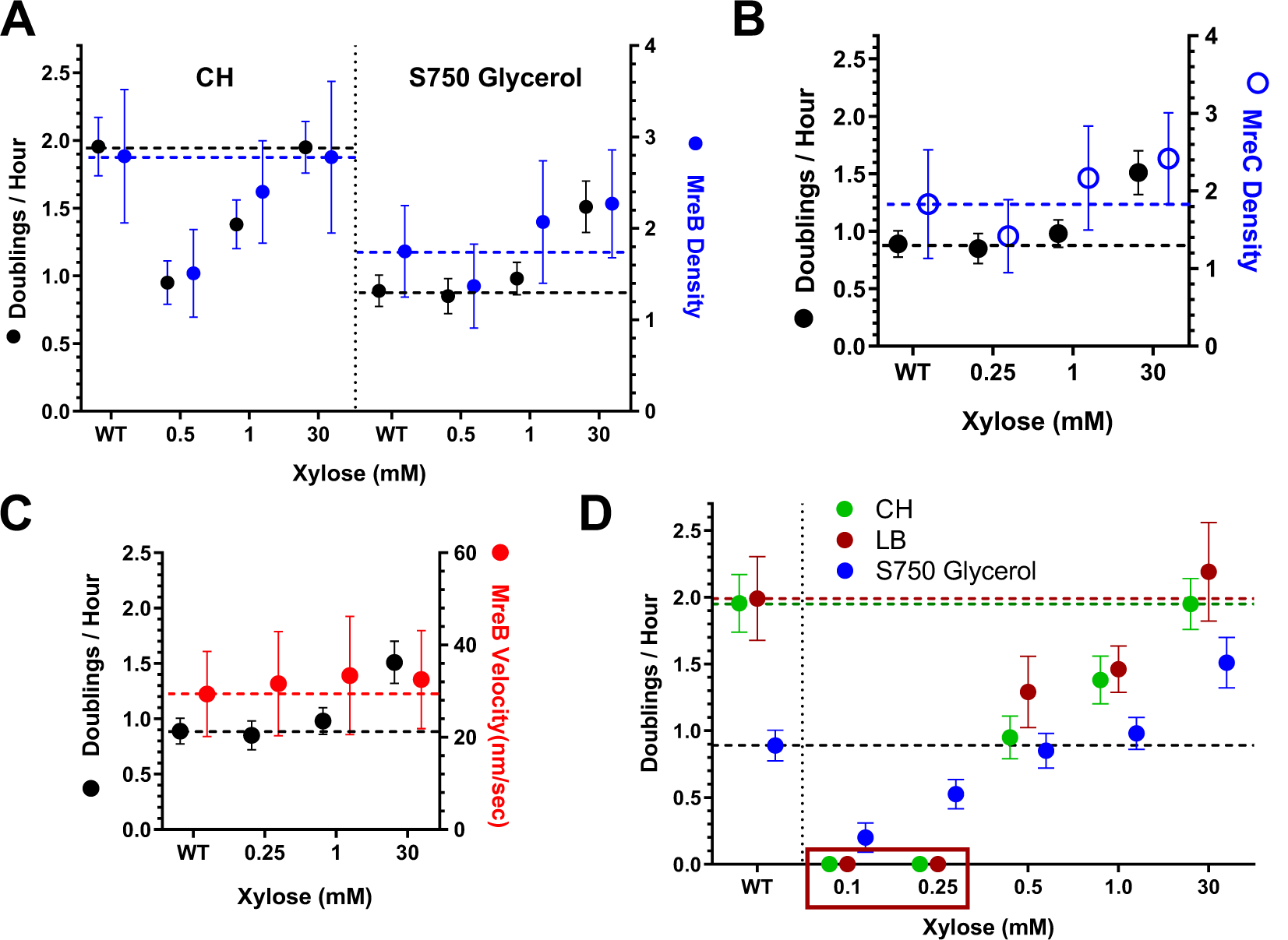
The density of directionally moving Rod complexes is regulated by *mur* activity. **(A) The density of MreB filaments can be titrated with MurAA expression -** Cells containing xylose inducible MurAA and MreB-mNeonGreen (bYS497) were grown in rich media (CH, *left*) or carbon limited media (S750 glycerol, *right*) at different *murAA* inductions, and their single-cell growth rate and directional MreB density determined. WT indicates cells with MreB-mNeonGreen with *murAA* under native control (bYS09). See also Movie S6. **(B) MreC density remains constant with MurAA induction -** Cells containing MreC-mNeonGreen and xylose inducible MurAA (bYS499) were grown with different inducer concentrations, and the directional MreC density determined. WT indicates cells containing MreC-mNeonGreen and MurAA under native control (bYS170). **(C) MreB velocity remains constant with MurAA induction -** Cells containing xylose inducible MurAA and MreB-mNeonGreen (bYS497) were grown in S750 glycerol with a range of MurAA inductions and the velocity of MreB filaments determined by particle tracking. **(D) Growth rate and MreB filament density can be titrated with different *murAA* inductions -** Cells with MurAA under xylose control (bYS365) were grown with different inducer concentrations, and their growth rates assayed in different media. Boxed region indicates the murAA inductions where cells do not grow, lysing instead. See also Fig. S2C. The growth rate, MreB filament density, and MreB velocity of cells with *murAA* under native control are illustrated by dashed lines in each figure.

When we repeated *murAA* titrations in cells growing in slow-growth carbon-limited media (S750 with glycerol as the carbon source) we observed a surprising effect: while decreased *murAA* inductions reduced both MreB density and cell growth, high *murAA* inductions not only increased MreB filament density 25% above that of WT, cells grew 30% faster than WT cells growing in the same media (Fig. 2D). While growth rate could be reduced by low *murAA* induction in all media tested, the growth acceleration only occurred in slow growth media, suggesting cells in rich media elongate at their maximal rate (Fig S2D). Thus, not only is both MreB filament density and overall growth rate regulated by the activity of the *mur* pathway, overexpression of *mur* genes somehow causes cells to grow faster than normal.

To test if MreB filament formation is regulated by interactions with Mur proteins, we tracked the single-molecule motions of the membrane-associated *mur* proteins MraY and MurG and the lipid II flippases MurJ and Amj (Meeske et al. 2015). In no case did we see any directional motions (Movie S7), indicating these proteins do not form stable interactions with the Rod complex. These results suggested lipid II signals to MreB via another manner. To determine if this signal arose from cytoplasmic or periplasmic facing lipid II, we observed MreB as we depleted both flippases, which should cause lipid II to accumulate inside the cell in the cytoplasmic orientation. Similar to previous *murG/murB* and *upps* depletions (Schirner et al. 2015), these depletions caused MreB filaments to depolymerize (Movie S8), indicating whatever signal controls MreB filaments arose from periplasmic oriented lipid II.

To determine what pathways sense periplasmic lipid II, we assayed the effects of chemical inhibitors on cell growth. One compound, staurosporine, reduced growth rate in minimal, but not rich media (Fig. 3A), suggesting signaling might be modulated through Hanks type kinases. *B. subtilis* possesses four serine/threonine kinases, and we assayed the growth of strains lacking each. The greatest reduction in growth occurred in cells lacking *prkC*, and a similar reduction occurred if we overexpressed the cognate phosphatase *prkC* (Fig. 3B).

**Fig. 3.**
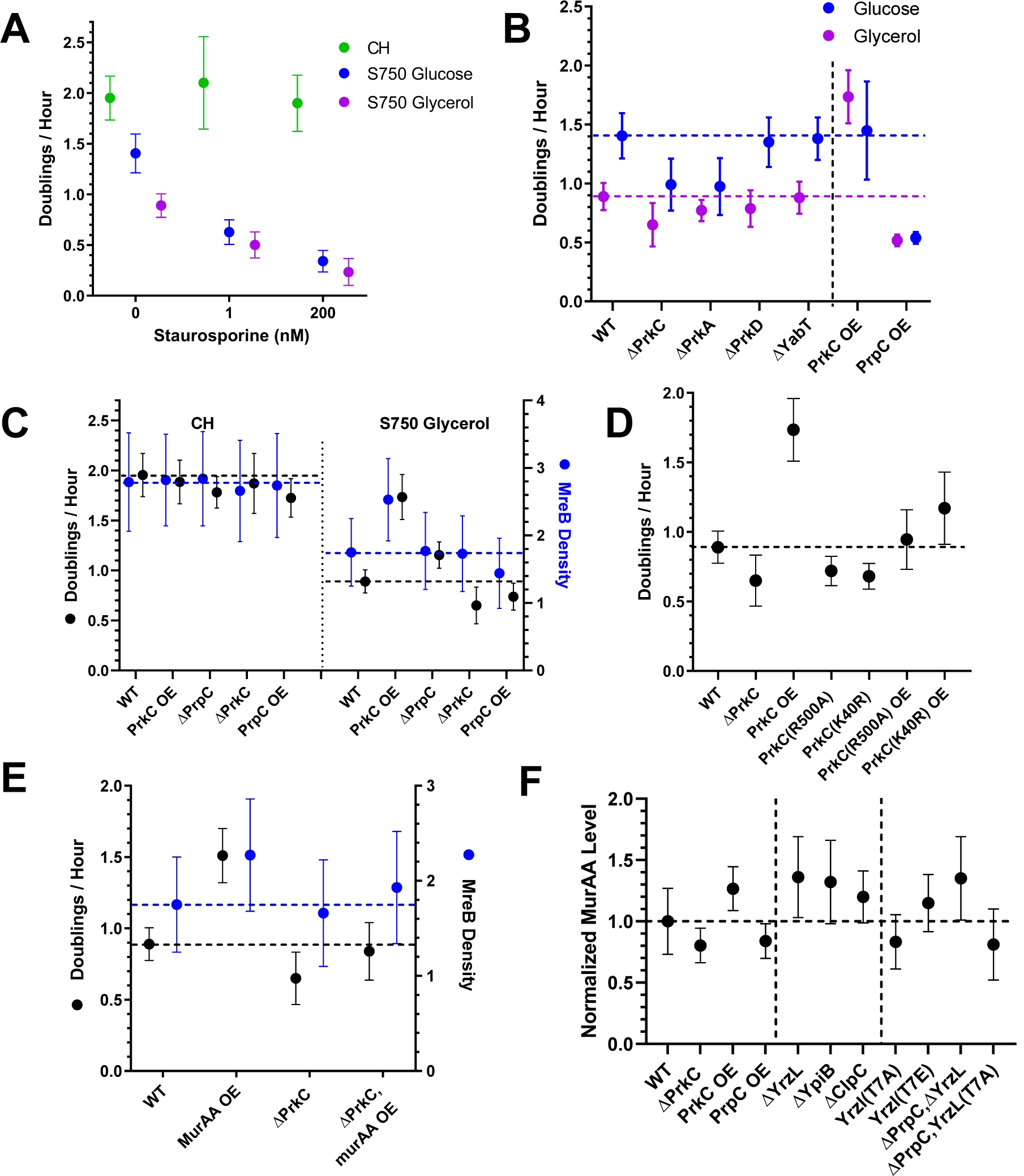
PrkC activity can both limit and overdrive cell growth. **(A) Staurosporine reduces the growth rate of cells in nutrient-limited media -** WT (PY79) cells were grown in different media at the indicated concentrations of staurosporine. **(B) Effects of kinases and phosphatases on cell growth** - The growth rate of cells in S750 glucose and glycerol was assayed in strains containing deletions of different Ser/Thr kinases (bYS542, bYS969, bYS970 and bYS971) and PrkC or PrpC overexpression (bYS730 and bYS545 with 30mM xylose). **(C) PrkC activity can limit and overdrive both growth rate and MreB filament density -** The growth rate and MreB filament density of strains modulating PrkC activity in nutrient-rich (*left*) and nutrient-limited media (*right*). PrkC and PrpC were overexpressed by growing bYS730 or bYS736, respectively, in 30mM xylose. *ΔprkC*: bYS736, *ΔprpC*: bYS732. **(D) PrkC’s lipid II binding and kinase activity are essential for growth acceleration** The growth rate of overexpressed PrkC mutants was assayed in S750 glycerol with 30mM xylose. PrkC(R500A) (bYS727) has a mutation in the PASTA domain. PrkC(K40R) (bYS729) has a mutation in the kinase catalytic domain. **(E) The *murAA* overexpression induced growth acceleration requires *prkC -*** The growth rate (in S750-glycerol) of Δ*prkC* cells overexpressing *murAA* (bYS619, induced with 30mM xylose) compared to cells only overexpressing *murAA* (bYS365, induced with 30mM xylose) or lacking *prkC*(bYS542). See also Fig. S3F-G. **(F) PrkC activity feeds back on *murAA* expression -** Integrated fluorescence of MurAA-mNeonGreen cells (WT, bYS397, grown in S750 glycerol), compared to cells overexpressing *prkC* (bYS1063, induced with 30mM xylose) or *prpC* (bYS1065, induced with 30mM xylose), Δ*prkC* (bYS893), Δ*yrzL* (bYS1069 and bYS1085), Δ*ypiB* (bYS1071), Δ*clpC* (bYS1067), YrzL T7E mutant (bYS1074) and YrzL T7A mutant (bYS1073 and bYS1084, induced with 1mM xylose). MurAA fluorescence is normalized to the level of WT MurAA-mNeonGreen cells. All data points represent fluorescence of >1000 cells. Dashed lines indicate growth rate and MreB filament density of WT (PY79) or control strains for each figure.

PrkC contains a cytoplasmic kinase domain and a periplasmic region containing Penicillin-binding And Serine/Threonine kinase Associated (PASTA) domains. The PASTA domains of PrkC and its homologs have been reported in various studies to bind to either lipid II or muropeptides (a moiety within lipid II) (Hardt et al. 2017; Kaur et al. 2019; Mir et al. 2011; Shah et al. 2008; Squeglia et al. 2011; Wamp, Rutter, Rismondo, Jennings, Möller, et al. 2020), thereby increasing the activity of the kinase domain. Mass spectrometry studies have indicated *prkC* and *prkC* levels are relatively constant across different growth conditions (Dion et al. 2019) (Fig. S3A).

Our next experiments revealed PrkC activity modulates the growth rate of cells in slow growth media: *prkC* overexpression caused cells to contain not only more MreB filaments, they also grew 53% faster than WT cells in the same media (Fig 3C, S3B). Likewise, Δ*prpC* cells showed more MreB filaments and grew 28% faster. Conversely, reducing PrkC activity by deleting *prkC* or overexpressing *prpC* reduced growth compared to WT and reduced MreB filament density. Similar to *murAA, prkC* overexpression increased the growth rate above that of WT in minimal, but not rich media (S3C). Growing cells in minimal media with different carbon sources revealed that PrkCs ability to tune growth has an upper limit: While cells growing slowly in poor carbon sources could have their growth reduced or accelerated by the respective removal or overexpression of *prkC*, cells growing much faster in glucose (an optimal carbon source) could only have their growth rate reduced, and not accelerated by *pkrC* overexpression beyond the already fast rate that WT cells exhibit in glucose containing media.

This growth acceleration required PrkC’s activities: cells encoding or overexpressing PrkC containing mutations that block pentapeptide binding (R500A) (Hardt et al. 2017) or abolish kinase activity (K40R) (Madec et al. 2002) caused no change relative to WT cells (Fig 3D). Furthermore, PrkC’s growth enhancing effect is downstream of MurAA, as the growth acceleration caused by *murAA* overexpression was eliminated in *ΔprkC* cells (Fig. 3E). Thus, under conditions of slow growth, suggest PrkC modulates both MreB filament density and overall growth rate by sensing the amount of lipid II in the periplasm and relaying this signal to the cytoplasm.

The PrkC mediated acceleration of growth posits a dilemma, as increased growth should require more lipid II, and thus require more *mur* activity. Recent work discovered a connection between PrkC and MurAA that might give this feedback: MurAA degradation is mediated by YrzL and YpiB, each of which increase the rate of MurAA proteolysis (Wamp, Rutter, Rismondo, Jennings, Moller, et al. 2020). Importantly, YrzL mediated degradation of MurAA is inhibited when it is phosphorylated by PrkC. To examine this finding in our context, we measured MurAA levels by fusing the native copy to mNeonGreen and quantitating cell fluorescence with widefield microscopy. Similar to observations by Wamp et al., *prkC* overexpression increased MurAA, and *prpC* overexpression reduced MurAA (Fig. 3F). Likewise, Δ*yrzL* and Δ*ypiB* cells showed increased MurAA. Importantly, blocking YrzL phosphorylation with a T7A mutation decreased MurAA levels, and the phosphomimetic mutation YrzL(T7E) increased MurAA. Accordingly, our proteomic data indicates that YrzL and YpiB levels decrease as cells are grown in increasingly rich media (Fig S3D). Thus, PrkC not only senses lipid II to adjust the rate of cell growth, it also feeds back on MurAA to modulate lipid II production.

How does PrkC modulate MreB filament number and overall growth rate? This is unlikely to occur via direct interactions with the Rod Complex, as single-molecule tracking of HaloTag-PrkC showed no directional motion, only diffusion within the membrane (Movie S9). Instead, we examined PrkC’s substrates. PrkC has been reported to phosphorylate numerous proteins (Ravikumar et al. 2018; Ravikumar et al. 2014), one of which is RodZ, a component of Rod complex, which we verified moves directionally using single-molecule tracking. Like MreC, an increasing number of RodZ molecules move directionally in faster growth media, while the rest diffuse along the membrane (Supplemental Text 2, Movie S10). RodZ is a transmembrane protein containing a periplasmic region that interacts with RodA/Pbp2a, and a cytoplasmic domain that binds to MreB (Bendezu et al. 2009; van den Ent et al. 2010). *E. coli* lacking RodZ or its cytoplasmic domain contain fewer MreB filaments, suggesting RodZ promotes filament formation via nucleation or stabilization (Hussain et al. 2018; Bratton et al. 2018). Phosphoproteomic studies have shown PrkC phosphorylates RodZ at serine 85, a residue in the linker adjacent to the MreB binding domain (Ravikumar et al. 2014; van den Ent et al. 2010).

We verified RodZ is phosphorylated at S85 by PrkC using PhosTag gel shifts (Fig. S4A). In line with PrkC phosphorylating RodZ in response to lipid II flux, RodZ phosphorylation increased as we titrated *murAA* expression (Fig 4A, Fig S4B). Likewise, RodZ phosphorylation scaled with the growth rate of cells growing in different amounts of different carbon sources (Fig. 4B).

**Fig. 4.**
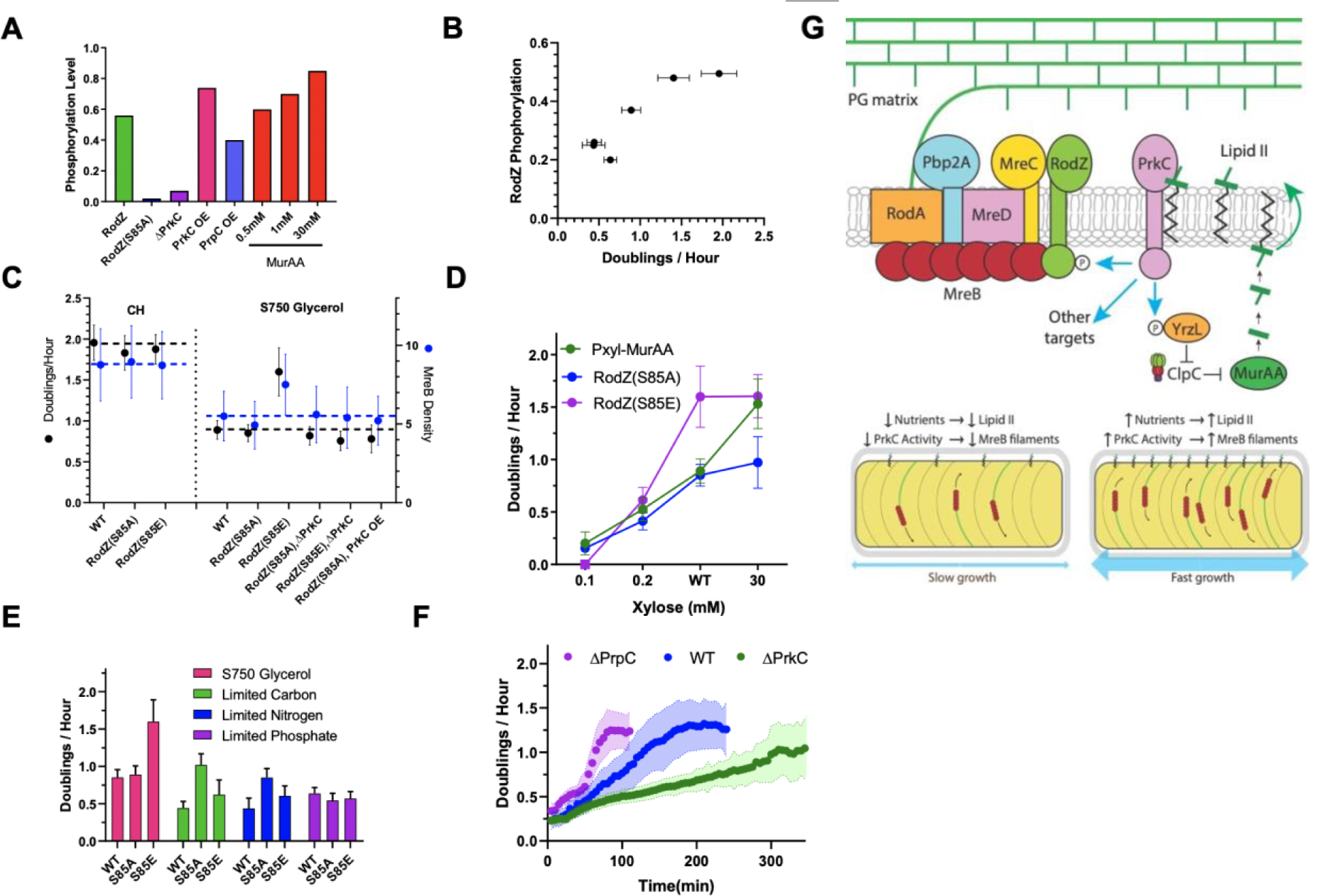
PrkC regulates cell growth via RodZ phosphorylation, creating a circuit helping cells grow across a range of nutrient conditions. **(A) RodZ phosphorylation depends on PrkC activity and MurAA induction -** PhosTag SDS PAGE was used to quantitate the fractional phosphorylation HaloTag-RodZ. All strains were grown in S750 glycerol. HaloTag-RodZ (bYS163), HaloTag-RodZ (S85A) (bYS165) and Δ*prkC* cells (bYS646) were labeled with 1 uM JF549 HaloTag. Overexpression of PrkC (bYS643) or PrpC (bYS644) in HaloTag-RodZ cells was induced by growth in 30mM xylose. HaloTag-RodZ cells containing xylose inducible MurAA (bYS642) were grown in xylose concentrations indicated. See also Fig.S4A-B. **(B) RodZ Phosphorylation scales with the growth rate of cells in different media -** HaloTag-RodZ (bYS163) was grown in different media, and the phosphorylation level of RodZ assayed using PhosTag SDS PAGE gels after cells were labeled with 1 uM JF549 HaloTag. **(C) Phosphorylation of RodZ controls MreB filament number and growth rate during slow growth -** *Left* - The growth rate and MreB density of cells grown in CH media, comparing cells with the phosphomimetic RodZ(S85E) mutation (bYS127), the phosphoablative mutation RodZ(S85A) (bYS125) to WT (PY79). *Right -* The growth rate and directional MreB density of cells grown in S750-glycerol, including the same RodZ mutants, *ΔprkC* (bYS542), or overexpressing *prkC* (bYS539 induced with 30mM xylose). **(D) PrkC mediated RodZ phosphorylation allows cells to survive across a range of lipid II levels -** The growth rate of WT RodZ(bYS365), RodZ(S85A)(bYS631), and RodZ(S85E)(bYS633) cells (grown in S750-glycerol) as MurAA was induced at different levels. At low MurAA inductions (0.1mM xylose), RodZ(S85E) cells stopped growing and underwent lysis. See also Fig. S4D-E. **(E) RodZ phosphorylation gives a growth advantage in excess nutrients, but a detriment when nutrients are limiting -** Growth rates of WT (PY79), RodZ(S85A)(bYS125), and RodZ(S85E)(bYS127) cells assayed in either S750-glycerol or the same media but where carbon (glycerol), nitrogen (glutamate) or phosphate (KH_2_PO_4_) were reduced by 50-fold. **(F) PrkC modulates the rate cells adapt to increased nutrients -** The growth rate of WT, Δ*prkC* (bYS542), and Δ*prpC* (bYS543) cells was assayed as they were shifted from growth in S750 glycerol (0.02%) to growth in S750 glycerol (1%). **(G) Model for regulation of cell growth by PrkC** *Top* - Schematic of PrkC mediated control of cell growth by lipid II. PrkC, by binding to lipid II, senses the amount, or flux, of cell wall precursors. Activated PrkC phosphorylates RodZ, YrzL, and other cellular targets. Phosphorylated RodZ then increases the amount of MreB filaments (either by nucleation or stabilization), which then activate the enzymes within the rod complex. PrkC also phosphorylates Yrzl, which increases the amount of MurAA by preventing its degradation by ClpC, thus increasing lipid II levels. *Bottom Left* - When nutrients are limiting, there is low rate of lipid II, and thus PrkC is relatively inactive. This causes RodZ to be mostly unphosphorylated, resulting in only a few MreB filaments per cell. *Bottom right* - As external nutrients increase, lipid II abundance increases, which leads to increased PrkC activity. This results in an increased phosphorylation of RodZ, which increases the number of MreB filaments in the cell, causing it to elongate faster.

To test if RodZ phosphorylation itself modulates cell growth, we created strains where S85 was mutated to alanine (S85A, mimicking dephosphorylation) or glutamic acid (S85E, mimicking phosphorylation). Strikingly, cells containing RodZ(S85E) not only showed more MreB filaments (Movie S11), they grew 66% faster than WT cells in the same media, close to the rate of WT cells in glucose. In glycerol, cells containing RodZ(S85A) showed a slightly reduced growth rate (6%) (Fig. 4C). The effects of RodZ phosphorylation are epistatic to MurAA and PrkC, as RodZ(S85A) abolished the faster growth of both *murAA* (Fig. 4D) and *prkC* overexpression (Fig. 4C). However, without *prkC*, RodZ (S85E) is not sufficient to increase growth rate, suggesting that PrkC must also phosphorylate other proteins in order for cells to achieve accelerated growth.

PrkC’s modulation of growth could serve different functions; First, it could help cells adapt to lipid II fluctuations, such as exposure to lipid II inhibiting antibiotics. Indeed, Δ*prkC* cells have growth defects relative to both WT and Δ*prpC* cells grown in the presence of sub-MIC levels of fosfomycin (Fig. S4D-F). More broadly, given PrkC phosphorylates hundreds of proteins involved in diverse essential processes (Libby, Goss, and Dworkin 2015; Libby, Reuveni, and Dworkin 2019; Pompeo et al. 2015; Shah et al. 2008), PrkC might act to simultaneously tune their overall activities in response to nutrient conditions. If so, decoupling one PrkC regulated process, such as cell wall synthesis, from the others should result in fitness defects, as this would cause one biosynthetic process to proceed too fast or slow relative to the others. To test this, we used the RodZ phospho-mutants to lock PrkC mediated control of PG synthesis into “on” or “off” states while we subjected cells at to extremes of *murAA* inductions or nutrient conditions. Indeed, cells containing RodZ (S85E) were unable to grow at low *murAA* inductions (Fig. 4C) and simply lysed, demonstrating that overactivation of PG synthesis is detrimental when the rest of the cell is growing slowly. Likewise, RodZ(S85A) cells grew slower than WT or RodZ(S85E) cells at high *murAA* induction, indicating cells cannot achieve the maximal possible growth rate when the PrkC regulation of PG synthesis is blocked. (Fig. 4C). Similar effects were observed when these strains were grown in media containing different concentrations of nutrients (Fig. 4D): In minimal media with saturating carbon (glycerol) or nitrogen (glutamate), “overactivated” RodZ(S85E) cells showed a growth advantage relative to WT and RodZ(S85A) cells. However, when carbon or nitrogen were limiting, cells grew much slower, often lysing. Likewise, “non-activatable” RodZ(S85A) cells were unable to attain the maximal growth rate of RodZ(S85E) in saturating nutrients, However, when nutrients were limiting, RodZ(S85A) cells showed a growth advantage relative to WT and RodZ(S85E) cells. We also observed the PrkC pathway is involved in how fast *B. subtilis* adapts to nutrient shifts. When shifting from low carbon media to carbon-rich media, Δ*prkC* cells were slower to adapt than WT cells, and conversely, Δ*prpC* cells adapt faster than both WT and Δ*prkC* cells (Fig. 4E).

Together, our data demonstrate that, rather than maximizing growth solely on limiting metabolites under conditions of slow growth, *B. subtilis* modulates both its amount of active cell wall synthesis enzymes and the overall rate of cell growth by measuring the flux of lipid II through cell wall synthesis. The cell’s metabolic state and PG synthetic activity are connected via PrkC, a kinase that phosphorylates multiple proteins when activated by lipid II. This signal is communicated to RodZ via phosphorylation, which then acts to increase the number of MreB filaments, thus increasing the number of directionally moving enzymes inserting material into the wall, thereby causing cells to elongate faster (Fig. 4F). Interestingly, PrkC also controls the rate of lipid II production via YrzL controlling MurAA stability (Wamp, Rutter, Rismondo, Jennings, Moller, et al. 2020, thus modulating the amount of the same molecular signal that controls its activity. This positive feedback might serve to increase the amount of PG precursors needed for increased Rod Complex activity as nutrients increase.

While this initial study focused on cell wall synthesis, PrkC and its homologs in other bacteria (Hardt et al. 2017; Cuenot et al. 2019; Wamp, Rutter, Rismondo, Jennings, Möller, et al. 2020) phosphorylate hundreds of proteins involved widely divergent cellular processes (Pereira, Goss, and Dworkin 2011; Ravikumar et al. 2018; Ravikumar et al. 2014), as well as controls cell growth via modulation of (p)ppGpp synthesis (Libby, Reuveni, and Dworkin 2019). This suggests that PrkC might function as a cellular rheostat, tuning the activities of many different cellular processes in response to lipid II so that their activities run at equivalent rates across the range of nutrient conditions.

Our findings also have implications for our understanding of the limitations of bacterial growth. While it has been long held that the limiting nutrient determines their maximal growth rate, these experiments reveal that, under limiting nutrient conditions, *B. subtilis* still contains the potential to grow ∼1.5 faster when given regulatory pathways are activated (Towbin et al. 2017; Cheng et al. 2014). Along with recent work demonstrating carbon limited bacteria contain an excess of inactive ribosomes (Bosdriesz et al. 2015), our findings suggest that, rather than growing at the maximal possible rate, bacteria actively hold back some growth capacity in reserve, activating or deactivating this capacity to tune their growth in response to the nutrient state or presence of antibiotics.

## Acknowledgments

We would like to thank P. Levin, E. Libby, S. Hürlimann, and A. Florez for helpful advice and discussions, M. Dion for strains, G. Squyres for reading the manuscript, and L. Lavis for JF dyes. This work was funded by National Institutes of Health Grants DP2AI117923-01 to EG, as well as support from the Volkswagen Foundation. This work was supported by the NSF-Simons Center for Mathematical and Statistical Analysis of Biology at Harvard (award number #1764269) and the Harvard Quantitative Biology Initiative.

## Author contributions

EG and YS conceived of the project and wrote the paper. YS conducted all experiments.

## Data and materials availability

Data generated and analyzed during this study are presented in the manuscript or in the supplementary materials; custom written codes for data analysis are available at the following sites: https://bitbucket.org/garnerlab/sun2019

## Supplementary Materials

Materials and Methods

Supplementary Texts 1 and 2

Supplementary Figures S1-S4

Supplementary Tables S1-S2

Supplementary Movie Captions SM1-SM10

## Supplementary Figures and Legends

**Fig. S1.**
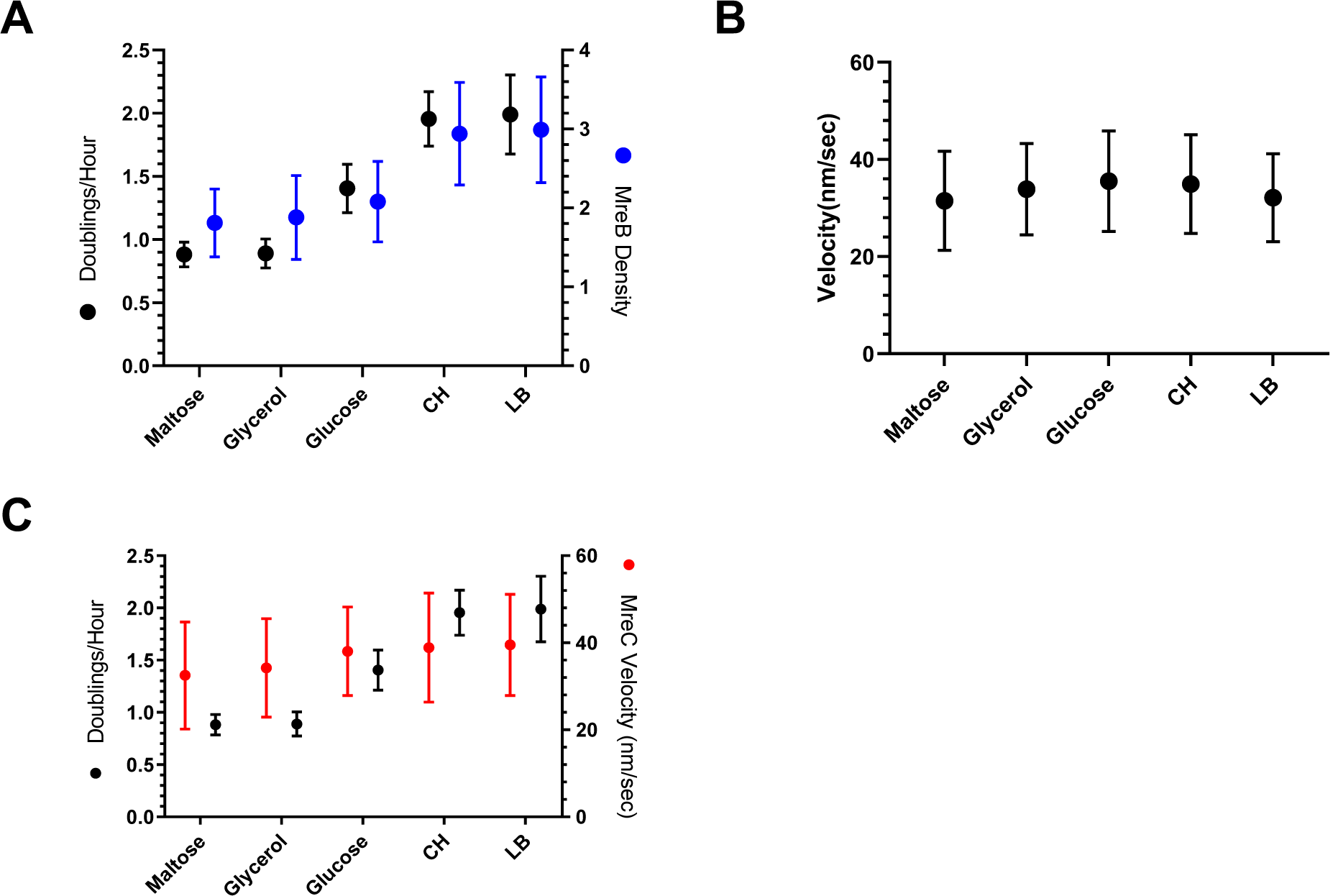
Quantitation of strains expressing MreB and Mbl fusions, and MreC velocity. (A) Growth rate and directional filament density and (B) velocity observed in cells expressing **<u>both</u>** MreB-sfGFP, Mbl-sfGFP(bMD203), taken in different media. (C) Velocity of MreC-mNeonGreen molecules (bYS170) in different media.

**Fig. S2.**
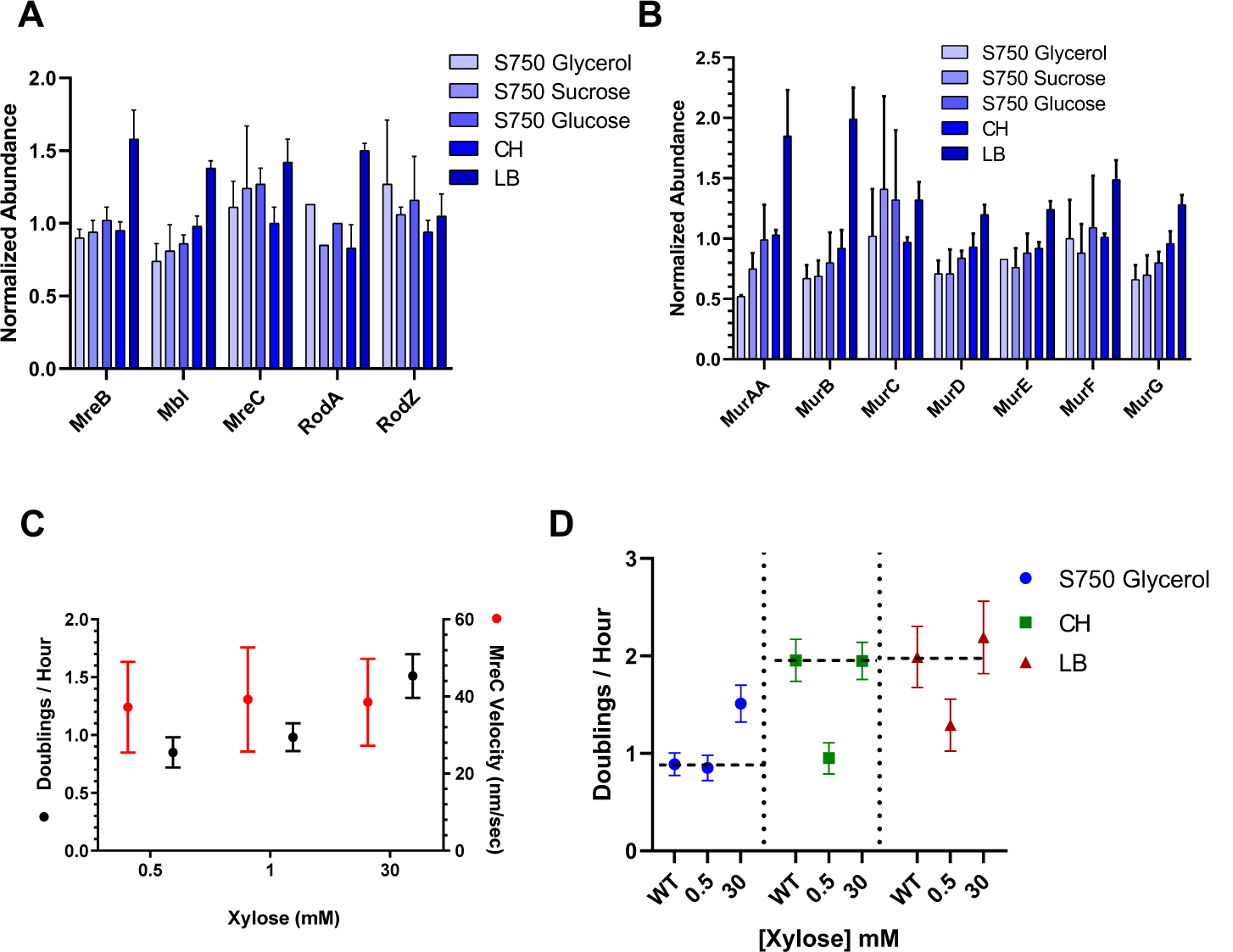
Relative abundance of Mur enzymes and effects of MurAA on MreB velocity and growth rate. (A) The relative abundance of Rod complex components in different media, as determined by proteomic mass spectrometry from (Dion et al. 2019). In this previous work, all abundances were normalized to the levels in WT(PY79) cells grown in CH. (B) The relative abundance of Mur enzymes in different media, again from (Dion et al. 2019). (C) MreC-mNeonGreen velocity during different inductions of MurAA(bYS499) grown in S750 glycerol. (D) Single-cell growth rates of WT(PY79) and Pxyl-MurAA (bYS365, with 0.5mM and 30mM xylose) in S750 glycerol, CH and LB.

**Fig. S3.**
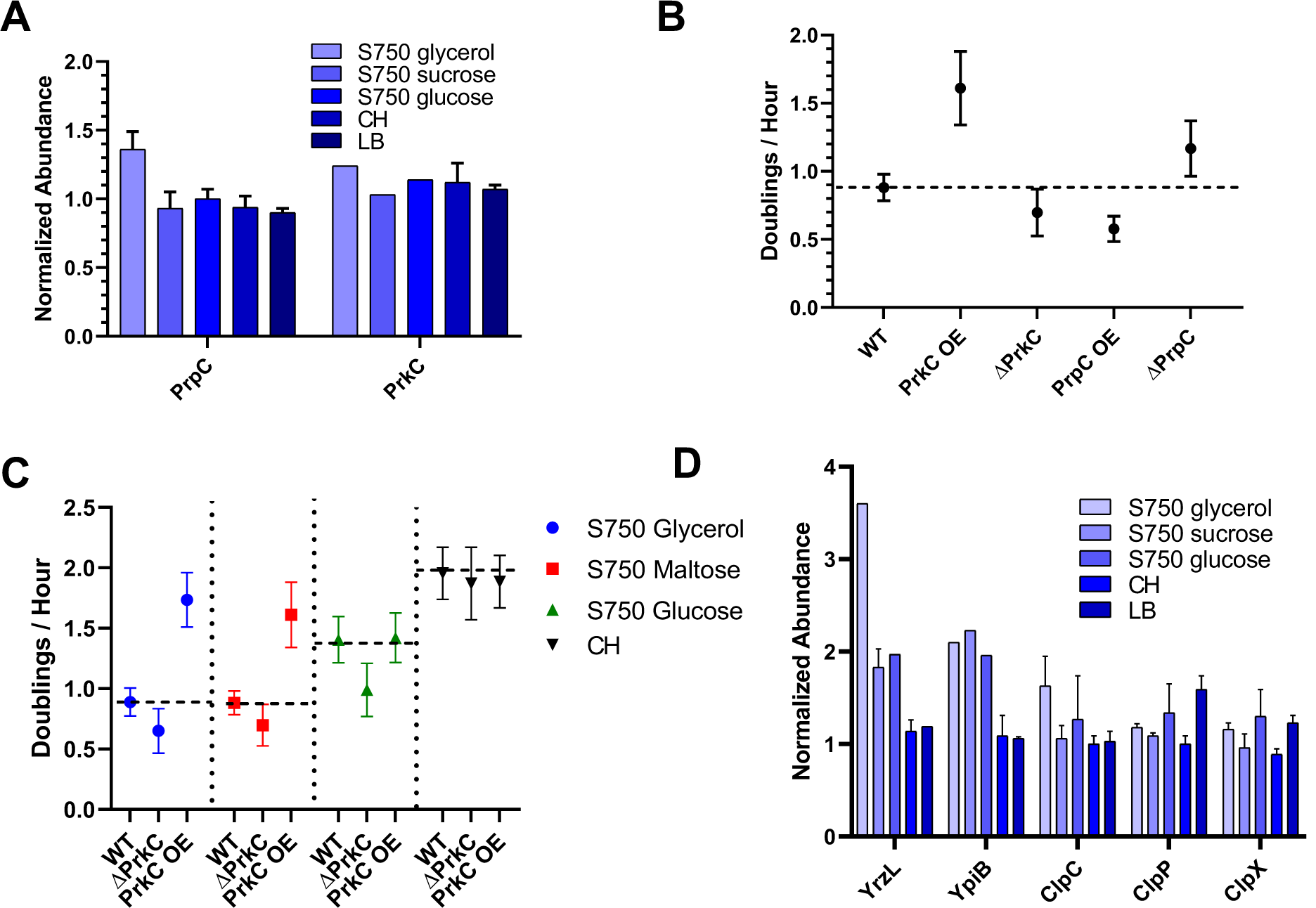
The Ser/Thr kinase PrkC regulates cell growth. (A) The relative abundance of PrpC and PrkC in WT (PY79) cells grown in different media. Data are from mass spectrometry studies in (Dion et al. 2019). (B) Growth rates of Δ*prkC* (bYS542), Δ*prpC* (bYS543), *prkC* overexpression (bYS539 with 30mM xylose), and *prpC* overexpression (bYS545 with 30mM xylose) of cells grown in S750 maltose. (C) Summary of growth rates of WT (PY79), Δ*prkC* (bYS542), and *prkC* overexpression (bYS539 with 30mM xylose) in S750 glycerol, maltose, glucose and CH. (D) The relative abundance of YrzL, YpiB, and ClpC/P/X protease in WT(PY79) in different media (Dion et al. 2019), showing the abundance of YrzL and YpiB decrease when cells are growing faster in richer media.

**Fig. S4.**
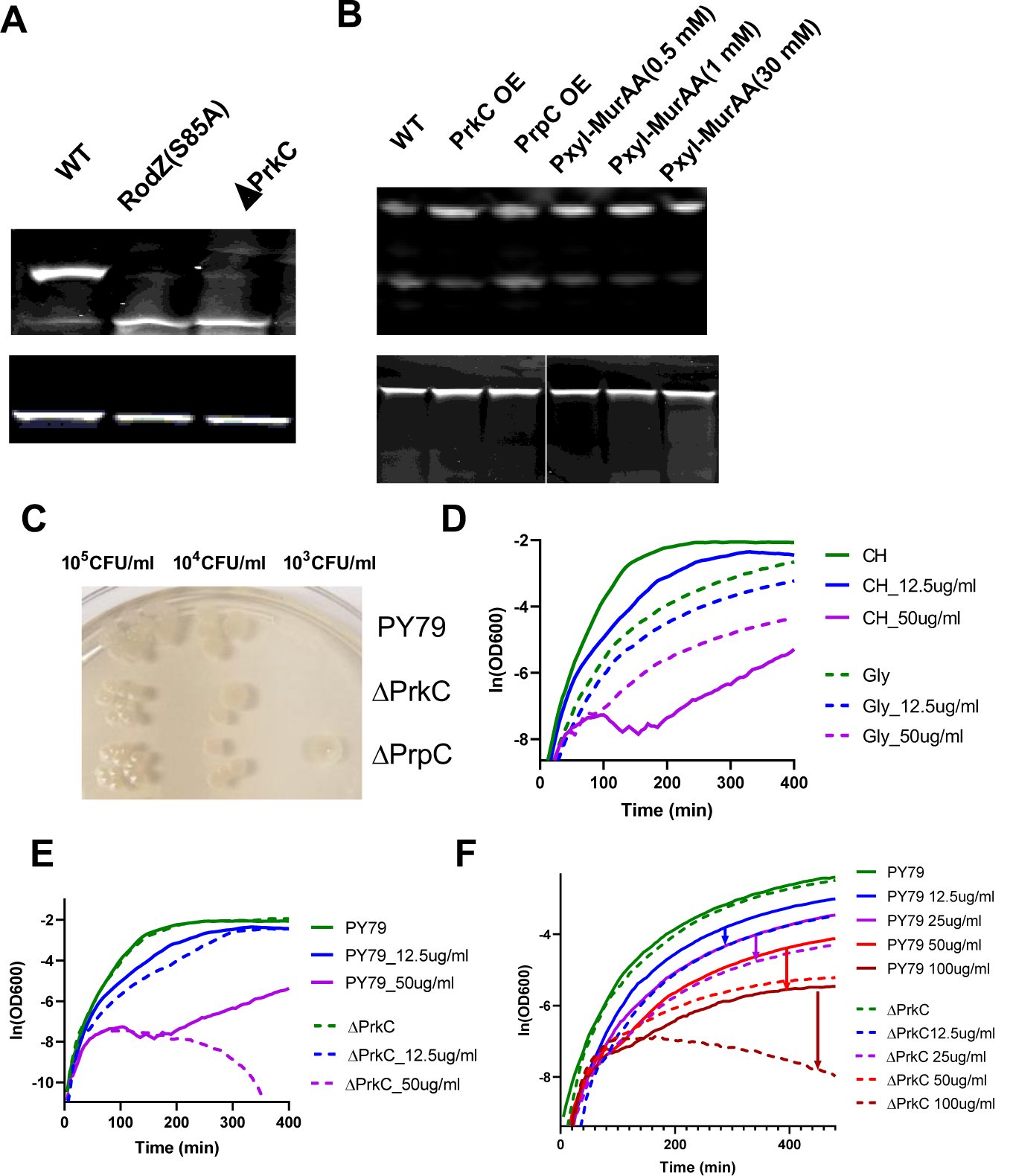
Phosphorylation of HaloTag-RodZ and effects of PrkC on fosfomycin sensitivity. (A) PhosTag SDS-PAGE gel shift of HaloTag-RodZ fusions. Top: SDS-PAGE supplemented with Phos-tag causes a band shift with HaloTag-RodZ (bYS163), but not in RodZ(S85A) (bYS165) or Δ*prkC* (bYS646) cells. Bottom: Regular SDS PAGE gel showing no band shift is observed. (B) Top: PhosTag SDS-PAGE of HaloTag-RodZ in cells overexpressing *prkC* (bYS643, 30mM xylose), overexpressing *prpC*(bYS644, 30mM xylose) and with different levels of *murAA*(bYS642) induction. Bottom: regular SDS PAGE gel, showing no second band is observed. (C) Spot plating assay of PY79, Δ*prkC* (bYS542), and Δ*prpC* (bYS543) cells on LB agar plates containing 25 ug/ml fosfomycin. The ability of the cells to grow on plates was examined after 12-hour incubation at 37°C. (D) Bulk growth (assayed using a shaking plate reader) of WT (PY79) cells, with different concentrations of fosfomycin in CH and S750 glycerol, respectively. (E, F) Bulk growth curves (assayed using a shaking plate reader) of WT(PY79) and Δ*prkC* (bYS542) cells grown in CH or S750 glycerol with different concentrations of fosfomycin.

## Supplementary Movie Captions

**Movie S1: Imaging of MreB filaments in different media**

TIRF-SIM imaging of MreB-mNeonGreen (bYS09) of cells grown in different media, imaged using TIRF-SIM microscopy at 1 second intervals with 100ms exposures.

**Movie S2: Single-molecule imaging of HaloTag-MreB sparsely labeled with JF549**.

HaloTag-MreB (bYS40) cells were incubated with 50 pM of HaloTag-JF549 ligand in different media for 30 minutes, washed, and prepared as indicated in methods. Cells were imaged at 0.5 second intervals using TIRFM with 0,5s exposures.

**Movie S3: Imaging of MreC-mNeonGreen in different media**

TIRF imaging of cells expressing MreC-mNeonGreen from the native locus (bYS170) grown in different media. Cells were imaged at 1 second intervals using TIRFM with 1s exposures.

**Movie S4: Imaging of MreB filaments during a nutrient upshift**

TIRF-SIM imaging of MreB-mNeonGreen (bYS09) during a nutrient upshift (from S750 maltose to CH starting at time 0. Cells were imaged at 1 second intervals with TIRF-SIM with 100ms exposures.

**Movie S5: Imaging of MreB and Mbl filaments in when cells are grown in different media** TIRF-SIM imaging of MreB-msfGFP, Mbl-msfGFP (bMD203) in different media. The time interval between frames was 1 second with 100ms exposures.

**Movie S6: Imaging of MreB filaments with different levels of MurAA induction**.

TIRF-SIM imaging of MreB-mNeonGreen, Pxyl-MurAA (bYS497) with different levels of MurAA induction. Xylose concentrations used to induce MurAA are indicated in each panel. Cells were imaged at 1 second intervals with an exposure time of 100ms.

**Movie S7: Imaging of MreB filaments with MurJ and Amj depletion**.

All cells were grown in CH or S750 glycerol and imaged at 1 second intervals using TIRFM. With an exposure time of 0.3 seconds.

Column 1: MreB-mNeonGreen (bYS09) growing in S750 glycerol.

Column 2: MreB-mNeonGreen, Pxyl-MurJ, Pxyl-Amj (bYS453) cells induced with 0.5mM xylose growing in S750 glycerol.

**Movie S8: TIRF imaging shows no directional motions of single molecules of MraY, MurG, and MurJ**.

All cells were grown in CH and imaged at 1 second intervals using TIRFM with 0.3s exposures.

Column 1: Imaging of msfGFP-MurG (bMD514).

Column 2: Imaging of Pxyl-mNeonGreen-MraY (bYS453) induced with 1mM xylose. Column 3: Imaging of Pxyl-mNeonGreen-MurJ (bYS609) induced with 1mM xylose. Column 4: Imaging of Pxyl-mNeonGreen-Amj (bYS611) induced with 1mM xylose.

**Movie S9: TIRF Imaging of HaloTag-PrkC in CH and glycerol**

HaloTag-PrkC (bYS765) was incubated with 10uM of HaloTag-JF549 ligand for 30 minutes, washed, then prepared as detailed in methods. Cells were imaged using TIRFM at 1 second intervals with 0.5s exposures.

**Movie S10: TIRF Imaging of HaloTag-RodZ**

Cells were imaged using TIRFM at 1 second intervals using an exposure time of 0.5 seconds. Column 1: Imaging of cells with HaloTag-RodZ at the native locus (bYS163) in CH. Column 2: Imaging of cells with HaloTag-RodZ (bYS163) in S750 glycerol.

**Movie S11: TIRF Imaging of MreB with RodZ mutants**

MreB-mNeonGreen (bYS09), MreB-mNeonGreen with RodZ(S85A) (bYS748), and MreB-mNeonGreen with RodZ(S85E) (bYS749) cells were grown in S750 maltose and S750 glycerol respectively. All cells were imaged at 1 second intervals using TIRFM with 0.3s exposures.

## Supplementary Text 1

*Note – this section will be updated in a updated preprint as the cross data analysis between labs evolves*.

Previously, Billaudeau and colleagues conducted a study detialing how the number of directionally moving MreB filaments (measured by particle tracking) changed with growth rate (Billaudeau et al. 2017). They came to the opposite conclusion that we arrive to here, that as *B. subtilis* grows faster, the filament velocity increases, not the number of MreB filaments.

We are confident in our data: we see same results not only with different fluorescent fusions to both MreB and Mbl; we gain the same results 1) conducting single-molecule tracking of sparsely labeled HaloTag-MreB, and, importantly, when 2) calculating the density and velocity MreC-mNeonGreen fusions. However, we trust in the good faith and scientific ability of colleagues, and thus, are working to understand the differences between our two studies. We are exchanging both data, strains, and code so that we can determine if the discrepancies in the two studies arise from 1) the strains and fusions used, or 2) the different analysis pipelines.

During the interim, we can only speculate why these differences occur. We think they may arise from either source:

1. The difference may arise from the MreB fusions used, as we and others have noted differences in the structures different MreB fusions make in the cell. For example, some fusions show numerous small filaments, while other fusions show fewer, but much longer filaments.
2. The difference may also come from the method of analysis, as while Billaudeau used particle tracking, which is subject to the diffraction limit. In contrast, our approach uses temporal correlations that between adjacent pixels and thus is able to count moving particles even when their density exceeds the diffraction limit (Dion et al. 2019). However, to control for this, we conducted single-particle tracking of sparsely labeled MreB Halo-Tag fusions and again saw no change in MreB velocity in different growth media. Thus, we currently disfavor the idea that these differences come from analysis methods, and likely think they arise from the fusion used.

We will update this preprint as we conduct our cross-lab comparison.

## Supplementary Text 2

Upon initiating our study of RodZ, we noted the correct open reading frame (ORF) of *Bacillus subtilis rodZ*(*ymfM*) was previously overlooked (https://www.uniprot.org/uniprot/O31771). We verified this using SDS PAGE gels, where HaloTag-RodZ (bYS163) (which contains an N-terminal HaloTag fusion directly before the overlooked upstream start codon) resulted in a single band at 70KDa (Fig. S4A, bottom panel). This indicated the full length of *B. subtilis* RodZ should be 304 aa instead of 288aa in the original annotation, where the first helix H1 of RodZ was truncated. Therefore, the phosphorylation site on RodZ should be S85 instead of S69 (Ravikumar et al. 2014). Single-molecule imaging of RodZ gave further verification this was the functional protein, showing that only HaloTag-RodZ (bYS163) with the correct ORF moves directionally around the cell width (Movie S10), while the original annotation, which truncated the protein (bYS151) moves freely, diffusing within the membrane, likely due to the lost interaction between MreB and RodZ (van den Ent et al. 2010).

## Materials and Methods

### Culture growth

Unless otherwise noted, *B. subtilis* was grown in casein hydrolysate (CH) medium or S750 minimal media with different carbon sources. Where indicated, xylose or isopropyl thiogalactoside (IPTG) was added. Cells were grown at 37°C to an OD600 of ∼0.4 to 0.6. The default concentration of carbon source was 1%.

### General imaging conditions

All components were prewarmed to 37 °C including Matek dishes, coverslips, agarose pads, and media. Exposure times varied between 300ms and 1 second, as indicated. And the interval between frames was 1 sec. A 488nm laser line was used for the imaging of mNeonGreen, and a 561nm laser line was used for the imaging of JF549-Halo dye.

### Single-cell growth rate time-lapse experiments

2ul of cell culture was spotted on No. 1.5 glass-bottomed dishes (MatTek, MA) under 3% agarose pads made with growth media. Phase-contrast images were collected on a Nikon Ti microscope equipped with a Hamamatsu ORCA Flash4 CMOS camera with a Nikon Plan Apo λ 100×/1.4NA objective with exposure times of 200ms, and each pixel was 64 nm. Images were acquired from 30-60 fields of view every 2 minutes for a total of 2-4 hours using Nikon NI Elements. Analysis of phase-contrast time-lapse movies was performed using a custom-built package in MATLAB. For nutrient upshift analyses, first cells were grown in slow growth media to mid-log (OD_600_ of ∼0.4 to 0.6) and then imaged under 3% agarose pads with the fast growth media. Phase-contrast time-lapse movies were then segmented, and single cell growth rates were calculated based on the piece-wise linear fitting of the area of cells. The instantaneous rate of growth during shift was calculated by rolling window regression using a window of 15 min.

### Bulk growth rate measurements

Cultures were grown at 37 °C to an OD600 of ∼0.6. The cultures were diluted back to a calculated OD 600 of 0.05 (replicated in 2-5 wells for each culture) and their growth measured using an Epoch microplate spectrophotometer (BioTek, VT) at 37 °C with continuous shaking. The growth rates were calculated from OD600 measurements that were recorded every 5 min for 6 h∼10h.

### TIRF-SIM Imaging of MreB

Cells were prepared as described in “Culture growth.” Cells were placed under an agarose pad in a No. 1.5 glass-bottomed dish (MatTek, MA) for imaging. Images were collected on a GE OMX in TIRF-SIM mode, using an Edge 5.5 sCMOS camera (PCO AG, Germany) and a 60x objective. 100 msec exposures from a 488 nm diode laser were used for each rotation. Raw images were reconstructed using SoftWoRx (GE Healthcare, MA) software. The pixel size after reconstruction was 40 nm.

### MreB-mNeonGreen and MreC-mNeonGreen dynamics assayed by TIRFM

Cells were prepared as described in “Culture growth.” Images were collected on a Nikon Ti microscope equipped with a Hamamatsu ORCA Flash4 CMOS camera with a Nikon 100X NA 1.45 objective. The pixel size was 65 nm. Cells were imaged with TIRFM for 120 seconds, followed by a single phase-contrast image.

For the nutrient upshifts conducted on strains expressing MreB-mNeonGreen, cultures grown in slow growth media to mid-log (OD_600_ of ∼0.4 to 0.6) were concentrated by centrifugation at 4500g for 60 seconds. The cell pellet was resuspended in nutrient-rich media (CH). 2ul of cell culture was then spotted on an ethanol-cleaned Matek disk with a No. 1.5 glass coverslip, under a 3% agarose pad containing CH, placed into a 37 °C chamber immediate imaged. Time-lapse imaging of MreB-mNeonGreen was performed every minutes 15 minutes with 300ms exposures, and the time interval is 1 sec.

### Imaging MreB-HaloTag, RodZ-HaloTag and PrkC-HaloTag by TIRFM

MreB-HaloTag cells were incubated with 50 pM of HaloTag-JF549 ligand for 30 minutes during growth at 37 °C with rotation and then washed twice with 2 volumes of growth media for sparse labeling. Cells were imaged at 1 second intervals using TIRF microscopy. The exposure time was 0.5 seconds. RodZ-HaloTag and PrkC-HaloTag were incubated with 1uM of HaloTag-JF549 ligand for 30 minutes for complete labeling. Following this, cells were washed twice with 2 volumes of growth media before imaging.

### Analysis of the density of MreB filaments and MreC enzymes

The method used to quantitate the density of MreB filaments and MreC enzymes has been published and extensively characterized previously (Dion et al. 2019). Briefly, phase images were segmented, and the width and midline of each cell were calculated. Next, the fluorescence time-lapses were analyzed based on the segmentation mask of the phase image. Cell contours and dimensions were calculated using the Morphometrics software package. Then kymographs were generated for each row of pixels along the midline of the cell, and the time-lapse movie for each cell was converted into a single 2D image. To identify filaments in the kymograph, closed image contours were generated in the 2D image. For these contours, we calculated the total intensity, the centroid and orientation. Next, to identify cases where the same MreB filament appears in multiple sequential kymographs, each object in a given kymograph was linked to a corresponding object in the previous and following kymographs based on the above properties of the object, and these objects counted as one event. All of the image analyses were performed using custom MATLAB code.

### Analysis of the velocity of directional moving MreB and MreC

Particle tracking of MreB and MreC was performed using the software package FIJI (Schindelin et al. 2012) and the TrackMate plugin (Tinevez et al. 2017). First, the phase images of cells were segmented, and then a segmentation mask was used to reject the tracks outside of the cells. And only tracks that are longer than seven frames were used for calculations of particle velocity. Particle velocity for each track was calculated based on nonlinear least-squares fitting using MSD(t)=4Dt+(vt)_2_, where MSD is the mean squared displacement, t is the time interval, D is the diffusion coefficient, and v is velocity. The time interval used for the velocity analysis was 80% of the track length. Tracks were excluded if the R-squared for the fitting of log[MSD] versus log[t] was less than 0.95. Single-molecule trajectories were discarded if displacement was < 250 nm.

### PhosTag SDS PAGE of HaloTag RodZ

5ml cultures were grown to OD600∼0.6 then the culture was labeled with JF549-Halo dye. The final concentration of JF549-Halo was 250nM. Cells were washed twice thoroughly with wash buffer (50 mM Tris pH 7.5, 150 mM NaCl, 150 mM NaF, and 2mM sodium orthovanadate) to remove free dye from the culture. Cells were lysed in lysis buffer (50 mM Tris pH 7.5, 150 mM NaCl, 3 mM MgCl2, 0.5% NP-40, 5 μg/ml DNase I (NEB), 1x PhosStop phosphatase inhibitor cocktail (Roche), 1 mM PMSF, and 1 mg/ml lysozyme to digest the cell wall). 12.5% resolving gels for SDS-PAGE were cast with 50 μM Phos-Tag acrylamide (Wako, AAL-107), and 5% stacking gels.

### Epifluorescence measurements of MurAA

Cells were prepared as described in “Culture growth.” Epifluorescence and phase images were collected using a Nikon Eclipse Ti equipped with a Nikon Plan Apo λ ×100/1.4NA objective and an Andor Clara camera. The pixel size was 64.5 nm. Exposure time is 500ms.

### Spot Plating Assay for the determination of survival of *B. Subtilis*

Cells were prepared as described in “Culture growth.” Overnight culture in LB or S750 glycerol was grown to OD600∼0.6. 8⨯10^8^ CFU/ml was used as the conversion ratio of OD600 to CFU/ml. Then the culture was serially diluted to obtain 1⨯10^5^, 1⨯10^4^, and 1⨯10^3^ CFU/ml culture. Then 10 μl of 1⨯10^5^, 1⨯10^4^, and 1⨯10^3^ CFU/ml culture were plated onto the LB plate respectively. The fosfomycin concentrations on LB plates were 25ug/ml and 50ug/ml. Plates were incubated at 37°C overnight.

## Supplemental Tables

**Table S1:** *B. subtilis* strains used in this study

| Strain | Relevant genotype | Source |
| --- | --- | --- |
| PY79 | wild type *B. subtilis* | BGSCID 1A747 |
| bYS009 | *mreB-mNeonGreen* | (Hussain et al. 2018) |
| bYS040 | *mreB-Halo* | (Hussain et al. 2018) |
| bYS121 | *rodZ(Y48A)* | This Work |
| bYS123 | *rodZ(Y48A, F53A)* | This Work |
| bYS125 | *rodZ(S85A)* | This Work |
| bYS127 | *rodZ(S85E)* | This Work |
| bYS163 | *Halo-rodZ* | This Work |
| bYS165 | *Halo-rodZ(S85A)* | This Work |
| bYS167 | *Halo-rodZ(S85E)* | This Work |
| bYS170 | *mNeonGreen-mreC* | This Work |
| bYS365 | *Pxyl-murAA::spec* | This Work |
| bYS385 | *Pxyl-mraY::cat* | This Work |
| bYS395 | *murAA-mNeonGreen::cat* | This Work |
| bYS397 | *murAA-mNeonGreen::loxP* | This Work |
| bYS395 | *Pxyl-murJ::cat* | This Work |
| bYS397 | *Pxyl-amj::cat* | This Work |
| bYS399 | *Pxyl-murJ::cat, Pxyl-amj::cat* | This Work |
| bYS453 | *Pxyl-mNeonGreen-mraY::cat* | This Work |
| bYS455 | *Pxyl-mNeoGreenn-murJ::erm* | This Work |
| bYS457 | *Pxyl-mNeoGreenn-amj::erm* | This Work |
| bYS497 | *mreB-mNeonGreen, Pxyl-murAA::spec* | This Work |
| bYS500 | *mNeonGreen-mreC, Pxyl-murAA::spec* | This Work |
| bYS539 | *amyE::Pxyl-prkC::kan* | This Work |
| bYS541 | $\Delta prkC$(LoxP) | This Work |
| bYS543 | $\Delta prpC$(LoxP) | This Work |
| bYS545 | *amyE::Pxyl-prpC::kan* | This Work |
| bYS567 | $\Delta prpC$, *rodZ(S85A)* | This Work |
| bYS721 | *amyE::Pxyl-prkC(R500A)::kan* | This Work |
| bYS723 | *amyE::Pxyl-prkC(R500E)::kan* | This Work |
| bYS725 | *amyE::Pxyl-prkC(K47R)::kan* | This Work |
| bYS619 | $\Delta prkC$, *Pxyl-murAA::spec* | This Work |
| bYS631 | *rodZ(S85A)*, *Pxyl-murAA::spec* | This Work |
| bYS633 | *rodZ(S85E)*, *Pxyl-murAA::spec* | This Work |
| bYS734 | $\Delta prkC$, *Pxyl-mreC::erm* | This Work |
| bYS735 | $\Delta prpC$, *Pxyl-mreC::erm* | This Work |
| bYS736 | *mreB-mNeonGreen, $\Delta prkC$* | This Work |
| bYS738 | *mreB-mNeonGreen, ΔprpC* | This Work |
| bYS746 | *rodZ(S85A), Pxyl-mreC::erm* | This Work |
| bYS747 | *rodZ(S85E), Pxyl-mreC::erm* | This Work |
| bYS748 | *mreB-mNeonGreen, rodZ(S85A)* | This Work |
| bYS749 | *mreB-mNeonGreen, rodZ(S85E)* | This Work |
| bYS763 | *HaloTag-PrkC::cat* | This Work |
| bYS765 | *HaloTag-PrkC* | This Work |
| bYS642 | *HaloTag-RodZ, Pxyl-murAA::spec* | This Work |
| bYS646 | *HaloTag-rodZ, ΔprkC* | This Work |
| bYS647 | *HaloTag -rodZ, ΔprpC* | This Work |
| bYS643 | *HaloTag -rodZ, amyE::Pxyl-prkC::kan* | This Work |
| bYS644 | *HaloTag -rodZ, amyE::Pxyl-prpC::erm* | This Work |
| bYS730 | *mreB-mNeonGreen, amyE::Pxyl-prkC::kan* | This Work |
| bYS732 | *mreB-mNeonGreen, amyE::Pxyl-prpC::kan* | This Work |
| bYS358 | *mreB-mNeonGreen, Pxyl-murG::cat* | This Work |
| bYS512 | *mreB-mNeonGreen, Pxyl-mraY::cat* | This Work |
| bYS670 | *mreB-mNeonGreen, Pxyl-murJ::cat, Pxyl-amj::cat* | This Work |
| bYS925 | *amyE::Phyperspank-ptpZ::erm* | This Work |
| bYS969 | *ΔprkA::cat* | This Work |
| bYS970 | *ΔprkD::cat* | This Work |
| bYS971 | *ΔyabT::cat* | This Work |
| bYS395 | *murAA-mNeonGreen::cat* | This Work |
| bYS397 | *murAA-mNeonGreen* | This Work |
| bYS1057 | *ΔyrzL::cat* | This Work |
| bYS1059 | *ΔypiB::cat* | This Work |
| bYS1061 | *AmyE::Pxyl-yrzL(T7A)::erm* | This Work |
| bYS1053 | *ΔclpC::cat* | This Work |
| bYS1055 | *ΔclpP::cat* | This Work |
| bYS893 | *murAA-mNeonGreen, ΔprkC* | This Work |
| bYS894 | *murAA-mNeonGreen, ΔprpC* | This Work |
| bYS895 | *murAA-mNeonGreen, AmyE::Phyper-prkC::erm* | This Work |
| bYS896 | *murAA-mNeonGreen, AmyE::Phyper-prpC::erm* | This Work |
| bYS1063 | *murAA-mNeonGreen, ΔclpC::cat* | This Work |
| bYS1065 | *murAA-mNeonGreen, ΔyrzL::cat* | This Work |
| bYS1067 | *murAA-mNeonGreen, ΔypiB::cat* | This Work |
| bYS1069 | *murAA-mNeonGreen, AmyE::Pxyl-yrzL(T7A)::erm* | This Work |
| bYS1070 | *murAA-mNeonGreen, ΔprpC, ΔyrzL::CAT* | This Work |
| bYS1071 | *murAA-mNeonGreen, ΔprpC, AmyE::Pxyl-yrzL(T7A)::erm* | This Work |
| bYS1072 | *murAA-mNeonGreen, ΔyrzL::cat, AmyE::Pxyl-yrzL(T7A)::erm* | This Work |
| bMD88 | *Pxyl-mreC::erm* | (Schirner et al. 2015) |
| bMD203 | *mreB-msfGFP, msfGFP-mbl::spec* | This Work |
| bMD514 | *sfGFP-40aa-murG::cat* | This Work |
| bMK169 | *Pxyl-rodZ::kan* | This Work |

**Table S2.**
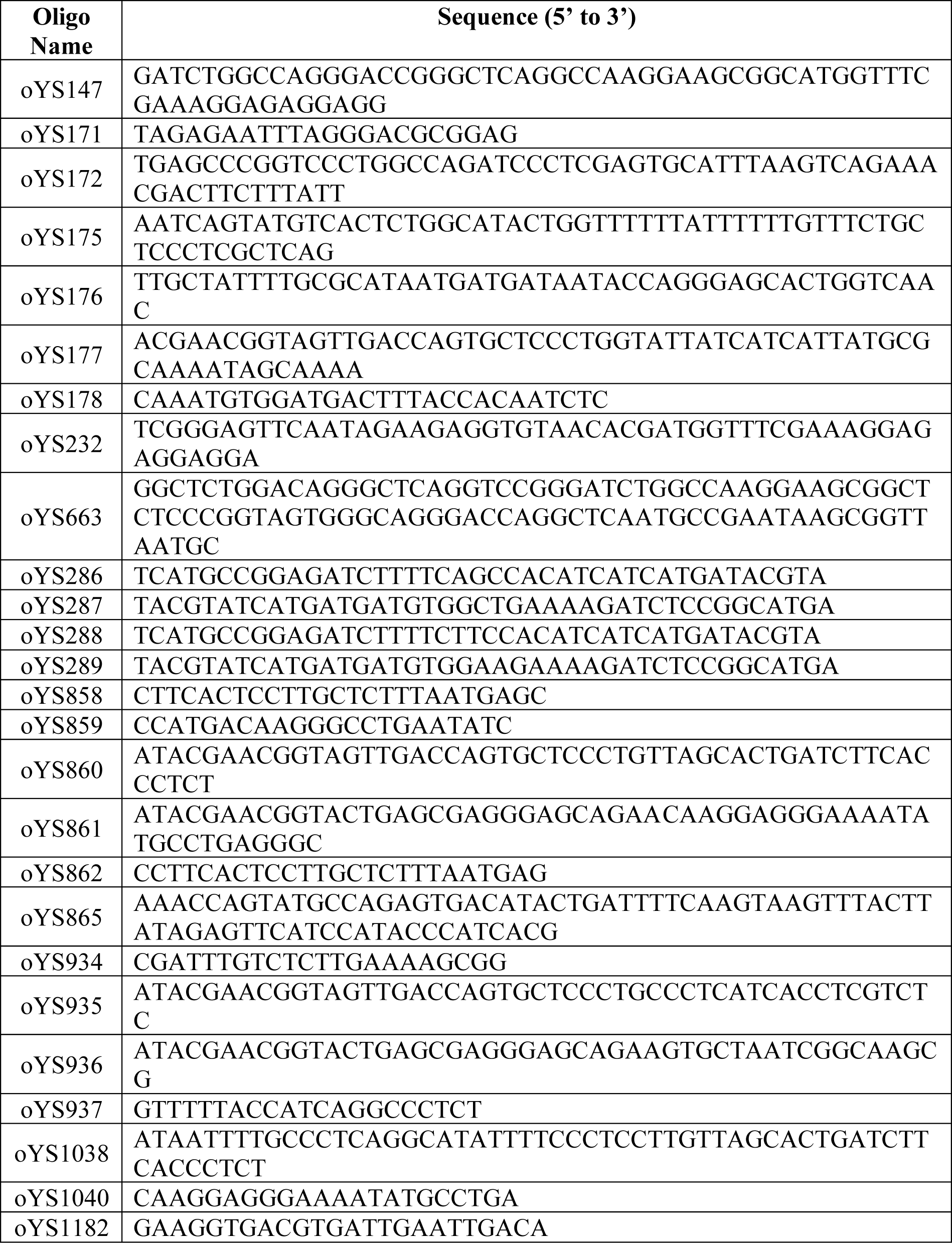
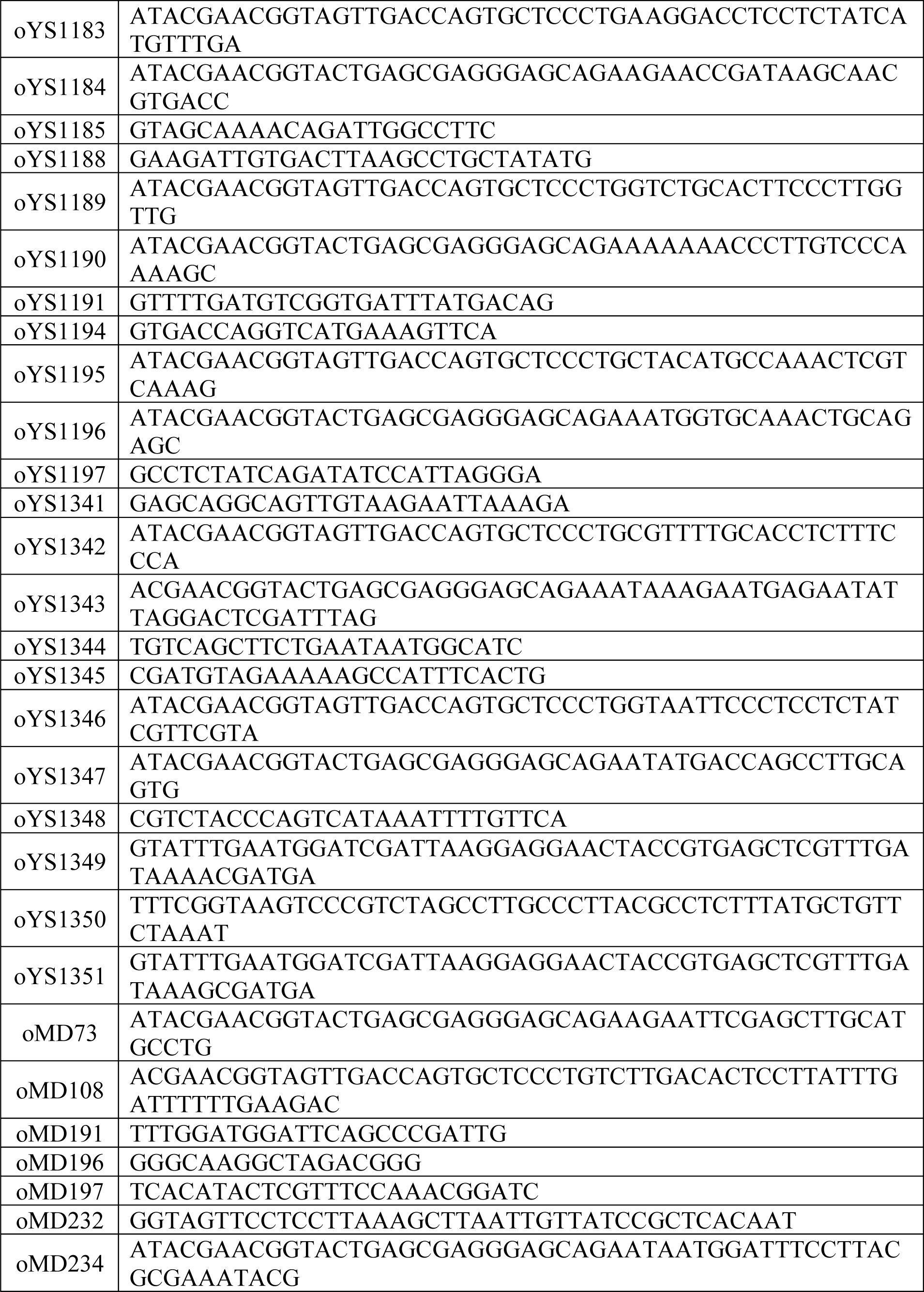
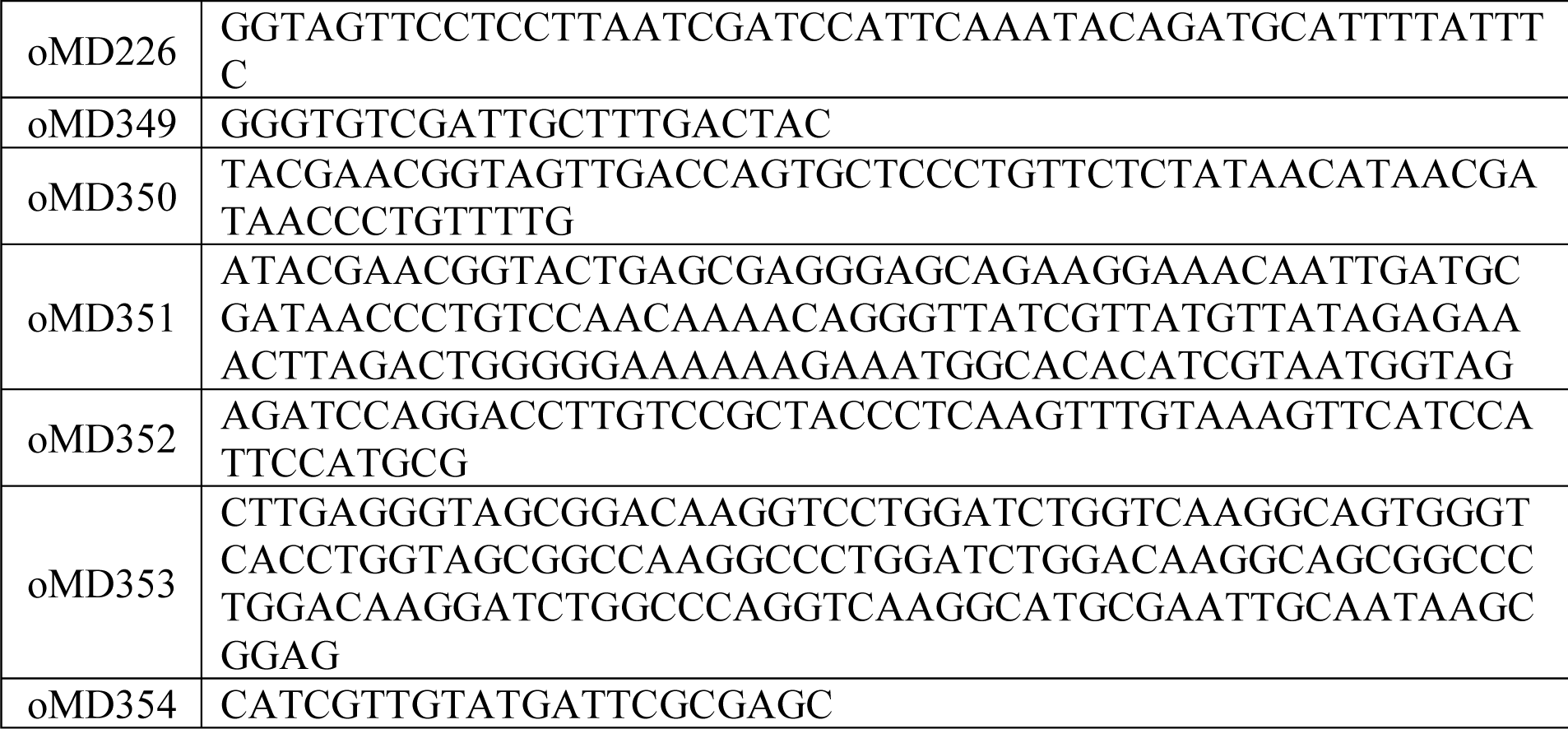
Oligonucleotides used in this study

## Strain Construction

**bYS170** harboring *mreC-mNeonGreen* was generated by transforming bMD88 (Schirner et al. 2015) with a Gibson assembly consisting of three fragments: (1) PCR with primers oMD117 and oYS230 and PY79 template genomic DNA (containing the upstream of *mreC*); (2) PCR with primers oYS232 and oYS233 and gBlocks gene fragment containing mNeonGreen; (3) PCR with primers oYS663 and oMD116 and PY79 template genomic DNA. Counterselection was done in the presence of 30 mM xylose. Selection on LB plates for growth in the absence of xylose and erm antibiotics resulted in strain bYS170. The genotype was confirmed by PCR and sequencing.

**bMK169** harboring *pxyl-rodZ::kan* was generated upon transformation of PY79 with a Gibson assembly consisting of four fragments: (1) PCR with primers oZB36 and oZB62 and PY79 template genomic DNA (containing the upstream or *rodZ*); (2) PCR with primers oJM028 and oJM029 and kanamycin-resistance cassette *loxP-kan-loxP* (amplified from pWX467 [a gift from D. Rudner]); (3) PCR with primers oMD73 and oMD226 and pDR150 template [gift of D. Rudner] (containing the *xylR* gene, and the *PxylA* promoter with an optimized ribosomal binding sequence; and (4) PCR with primers oMK106 and oZB44 and PY79 template genomic DNA (containing the *rodZ* coding region).

**bYS125** harboring *rodZ(S85A)* was generated by transforming bMK169 with a Gibson assembly consisting of two fragments: 1) PCR with primers oZB36 and oYS286 and PY79 template genomic DNA (containing the upstream of *rodZ* locus); (2) PCR with primers oYS287 and oZB44 and PY79 template genomic DNA (containing the downstream of *rodZ* locus). Selection on LB plates for growth in the absence of xylose and kan antibiotics resulted in strain bYS125. Counterselection was done in the presence of 30 mM xylose. The genotype was confirmed by PCR and sequencing.

**bYS127** harboring *rodZ(S85E)* was generated by transforming bMK169 with a Gibson assembly consisting of two fragments: 1) PCR with primers oZB36 and oYS288 and PY79 template genomic DNA (containing the upstream of *rodZ* locus); (2) PCR with primers oYS289 and oZB44 and PY79 template genomic DNA (containing the downstream of *rodZ* locus). Selection on LB plates for growth in the absence of xylose and kan antibiotics resulted in strain bYS127. Counterselection was done in the presence of 30 mM xylose. The genotype was confirmed by PCR and sequencing.

**bYS151** harboring NTD truncated *HaloTag-rodZ* was generated by transforming bMK169 with a Gibson assembly consisting of three fragments: 1) PCR with primers oZB36 and oYS290 and PY79 template genomic DNA (containing the upstream of *rodZ* locus); (2) PCR with primers oYS616 and oYS601 and Halo-tag template genomic DNA (3) PCR with primers oYS291 and oZB44 and PY79 template genomic DNA (containing the downstream of *rodZ* locus). Selection on LB plates for growth in the absence of xylose and kan antibiotics resulted in strain bYS151. Counterselection was done in the presence of 30 mM xylose. The genotype was confirmed by PCR and sequencing.

**bYS163** harboring *HaloTag-rodZ* was generated by transforming bMK169 with a Gibson assembly consisting of three fragments: 1) PCR with primers oZB36 and oYS290 and PY79 template genomic DNA (containing the upstream of *rodZ* locus); (2) PCR with primers oYS616 and oYS601 and Halo-tag template genomic DNA (3) PCR with primers oYS291 and oZB44 and PY79 template genomic DNA (containing the downstream of *rodZ* locus). Selection on LB plates for growth in the absence of xylose and kan antibiotics resulted in strain bYS163. Counterselection was done in the presence of 30 mM xylose. The genotype was confirmed by PCR and sequencing.

**bYS165** harboring *HaloTag-rodZ(S85A)* was generated by transforming bMK169 with a Gibson assembly consisting of two fragments: 1) PCR with primers oZB36 and oYS286 and bYS163 template genomic DNA(containing the upstream of *rodZ* locus and HaloTag); (2) PCR with primers oYS287 and oZB44 and PY79 template genomic DNA. Selection on LB plates for growth in the absence of xylose and kan antibiotics resulted in strain bYS165. Counterselection was done in the presence of 30 mM xylose. The genotype was confirmed by PCR and sequencing.

**bYS365** harboring *pxyl-murAA::spec* was generated upon transformation of PY79 with a four-piece Gibson assembly reaction, that contained the following PCR products. (1) PCR with primers oMD388 and oMD389 and PY79 template genomic DNA.; (2) PCR with primers oJM028 and oJM029 and the spectinomycin-resistance cassette *loxP-spec-loxP*(amplified from pWX467 [gift of D. Rudner]); (3) a fragment containing the *xylR* gene, and the *PxylA* promoter with an optimized ribosomal binding sequence (amplified from pDR150 using primers oMD73 and oMD226); and (4) a 1271 bp fragment containing the *murAA* coding region (amplified from PY79 genomic DNA using primers oMD394 and oMD395).

**bYS395** harboring *murAA-mNeonGreen::cat* was generated upon transformation of PY79 with a Gibson assembly consisting of three fragments: (1) PCR with primers oYS171 and oYS172 and PY79 template genomic DNA (containing the upstream of *murAA* locus); (2) PCR with primers oYS147 and oYS865 and gBlock gene fragment containing mNeonGreen template DNA (3) PCR with primers oYS175 and oYS176 and chloramphenicol-resistance cassette *loxP-cat-loxP* template (amplified from pWX465[a gift from D. Rudner]); and (4) PCR with primers oYS177 and oYS178 and PY79 template genomic DNA.

**bYS397** harboring *murAA-mNeonGreen::loxP*. The *loxP-cat-loxP* antibiotics cassette was subsequently looped out using a cre-expressing plasmid pDR244 (Koo et al. 2017).

**bYS537** harboring *ΔprkC::cat* is a *prkC* deletion strain. It was generated upon transformation of PY79 with a three-piece Gibson assembly reaction. 1) PCR with primers oYS859 and oYS860 and PY79 template genomic DNA (containing the upstream of *prkC* locus); (2) PCR with primers oJM028 and oJM029 and *loxP-cat-loxP* template (amplified from pWX465); (3) PCR with primers oYS861 and oYS862 and PY79 template genomic DNA (containing the downstream of *prkC* locus).

**bYS542** harboring *ΔprkC* is a markerless *prkC* deletion strain. The *loxP-cat-loxP* antibiotics cassette of bYS537 was looped out using a cre-expressing plasmid pDR244.

**bYS539** harboring *amyE::Pxyl-prkC::kan* was generated upon transformation of PY79 with a four-piece Gibson assembly reaction, that contained the following PCR products. (1) PCR with primers oMD191 and oMD108 and PY79 template genomic DNA; (2) PCR with primers oJM028 and oJM029 and *loxP-kan-loxP* template (amplified from pWX470); (3) PCR with primers oMD73 and oMD226 and pDR150 [gift of D. Rudner] template DNA (containing the *xylR* gene, and the *PxylA* promoter with an optimized ribosomal binding); (4) PCR with primers oYS1014 and oYS1015 and PY79 template genomic DNA; and (5) PCR with primers oMD196 and oMD197 and PY79 template genomic DNA.

**bYS543** harboring *ΔprpC::erm* is a prpC deletion strain. It was generated upon transformation of PY79 with a three-piece Gibson assembly reaction that contained the following PCR products. 1) PCR with primers oYS934 and oYS935 and PY79 template genomic DNA (containing the upstream of *prpC* locus); (2) PCR with primers oJM028 and oJM029 and *loxP-erm-loxP* template (amplified from pWX467 [gift of D. Rudner]); (3) PCR with primers oYS936 and oYS937 and PY79 template genomic DNA. The *loxP-erm-loxP* antibiotics cassette was subsequently looped out using a cre-expressing plasmid pDR244.

**bYS545** harboring *amyE::Pxyl-prpC::erm* was generated upon transformation of PY79 with a four-piece Gibson assembly reaction, that contained the following PCR products: (1) A 1228 bp fragment containing sequence upstream of the *amyE* locus (amplified from PY79 genomic DNA using primers oMD191 and oMD108); (2) the 1673 bp erythromycin-resistance cassette *loxP-erm-loxP* (amplified from pWX467 [a gift from D. Rudner] using primers oJM028 and oJM029); (3) a 1532 bp fragment containing the *xylR* gene, and the *PxylA* promoter with an optimized ribosomal binding sequence (amplified from pDR150 [a gift from D. Rudner] using primers oMD73 and oMD226); (4) a 906 bp fragment containing the *PrpC* (amplified from PY79 genomic DNA using primers oYS1014 and oYS1015); and (5) a 1216 bp fragment containing the *amyE* terminator, and sequence downstream of the *amyE* locus (amplified from PY79 genomic DNA using primers oMD196 and oMD197).

**bYS549** harboring *amyE::Phyper-PrkC::erm* was generated upon transformation of PY79 with a five-piece Gibson assembly reaction, that contained the following PCR products. (1) PCR with primers oMD191 and oMD108and PY79 template genomic DNA; (2) PCR with primers oJM028 and oJM029 and *loxP-cat-loxP* template (amplified from pWX465); (3) PCR with primers oMD232 and oMD234 and pDR111 [gift of D. Rudner] template genomic DNA (containing the lacI gene, and the Phyperspank promoter with an optimized ribosomal binding sequence) (4) PCR with primers oYS864 and oYS981 and PY79 template genomic DNA; and (5) PCR with primers oMD196 and oMD197 and PY79 template genomic DNA.

**bYS551** harboring *amyE::Phyper-PrpC::erm* was generated upon transformation of PY79 with a five-piece Gibson assembly reaction, that contained the following PCR products. (1) PCR with primers oMD191 and oMD108and PY79 template genomic DNA; (2) PCR with primers oJM028 and oJM029 and *loxP-cat-loxP* template (amplified from pWX465); (3) PCR with primers oMD232 and oMD234 and pDR111 [gift of D. Rudner] template genomic DNA (containing the lacI gene, and the Phyperspank promoter with an optimized ribosomal binding sequence) (4) PCR with primers oYS1014 and oYS1015 and PY79 template genomic DNA; and (5) PCR with primers oMD196 and oMD197 and PY79 template genomic DNA.

**bYS497** harboring *mreB-mNeonGreen, Pxyl-murAA::spec* was generated upon transformation of bYS09 with genomic DNA from bYS365.

**bYS500** harboring *mNeonGreen-mreC, Pxyl-murAA::spec* was generated upon transformation of bYS170 with genomic DNA from bYS365.

**bYS642** harboring *HaloTag-RodZ, Pxyl-murAA::spec* was generated upon transformation of bYS163 with genomic DNA from bYS365.

**bYS646** harboring *HaloTag-RodZ, prkC KO* was generated upon transformation of bYS163 with genomic DNA from gYS537. The *loxP-cat-loxP* antibiotics cassette was then looped out using a cre-expressing plasmid pDR244.

**bYS647** harboring *HaloTag -RodZ, prpC KO* was generated upon transformation of bYS163 with genomic DNA from bYS543. The *loxP-erm-loxP* antibiotics cassette was then looped out using a cre-expressing plasmid pDR244.

**bYS643** harboring *HaloTag -RodZ, amyE::Pxyl-prkC::kan* was generated upon transformation of bYS163 with genomic DNA from bYS539.

**bYS644** harboring *HaloTag -RodZ, amyE::Pxyl-prpC::erm* was generated upon transformation of bYS163 with genomic DNA from bYS545.

**bYS721** harboring *amyE::Pxyl-prkC(R500A)::kan* was generated upon transformation of PY79 with a four-piece Gibson assembly reaction, that contained the following PCR products. (1) PCR with primers oMD191 and oMD108 and PY79 template genomic DNA; (2) PCR with primers oJM028 and oJM029 and *loxP-kan-loxP* template (amplified from pWX470); (3) PCR with primers oMD73 and oMD226 and pDR150 template DNA (containing the *xylR* gene, and the *PxylA* promoter with an optimized ribosomal binding); (4) PCR with primers oYS1014 and oYS1015 and PY79 template genomic DNA; and (5) PCR with primers oMD196 and oMD197 and PY79 template genomic DNA.

**bYS725** harboring *amyE::Pxyl-prkC(K40R)::kan* was generated upon transformation of PY79 with a four-piece Gibson assembly reaction, that contained the following PCR products. (1) PCR with primers oMD191 and oMD108 and PY79 template genomic DNA; (2) PCR with primers oJM028 and oJM029 and *loxP-kan-loxP* template (pWX470); (3) PCR with primers oMD73 and oMD226 and pDR150 template DNA (containing the *xylR* gene, and the *PxylA* promoter with an optimized ribosomal binding); (4) PCR with primers oYS1014 and oYS1015 and PY79 template genomic DNA; and (5) PCR with primers oMD196 and oMD197 and PY79 template genomic DNA.

**bYS734** harboring *ΔprkC, Pxyl-mreC::erm* was generated upon transformation of bYS542 with genomic DNA from bMD88.

**bYS735** harboring *ΔprpC, Pxyl-mreC::erm* was generated upon transformation of bYS543 with genomic DNA from bMD88.

**bYS746** harboring *rodZ(S85A), Pxyl-mreC::erm* was generated upon transformation of bYS125 with genomic DNA from bMD88.

**bYS747** harboring *rodZ(S85E), Pxyl-mreC::erm* was generated upon transformation of bYS127 with genomic DNA from bMD88.

**bYS736** harboring *ΔprkC, mreB-mNeonGreen* was generated upon transformation of bYS734 with genomic DNA from bYS09. Counterselection was done in the presence of 30 mM xylose. Selection on LB plates for growth in the absence of xylose and erm antibiotics resulted in strain bYS736.

**bYS738** harboring *ΔprpC, mreB-mNeonGreen* was generated upon transformation of bYS735 with genomic DNA from bYS09. Counterselection was done in the presence of 30 mM xylose. Selection on LB plates for growth in the absence of xylose and erm antibiotics resulted in strain bYS738.

**bYS748** harboring *rodZ(S85A), mreB-mNeonGreen* was generated upon transformation of bYS746 with genomic DNA from bYS09. Counterselection was done in the presence of 30 mM xylose. Selection on LB plates for growth in the absence of xylose and erm antibiotics resulted in strain bYS748.

**bYS749** harboring *rodZ(S85E), mreB-mNeonGreen* was generated upon transformation of bYS747 with genomic DNA from bYS09. Counterselection was done in the presence of 30 mM xylose. Selection on LB plates for growth in the absence of xylose and erm antibiotics resulted in strain bYS749.

**bYS763** harboring *HaloTag-prkC::cat* was generated by transforming bYS542 with a Gibson assembly consisting of five fragments: 1) PCR with primers oMD464 and oYS1031 and PY79 template genomic DNA (containing the upstream of *prkC* locus); (2) PCR with primers oYS616 and oAB60 and Halo-tag template genomic DNA; (3) PCR with primers oYS1086 and oYS984 and PY79 template genomic DNA (4) PCR with primers oJM028 and oJM029 and *loxP-cat-loxP* template (amplified from pWX465); (5) PCR with primers oYS861 and oYS862 and PY79 template genomic DNA (containing the downstream of *prkC* locus).

**bYS765** harboring *HaloTag-prkC* is a markerless *HaloTag-prkC* strain. The *loxP-cat-loxP* antibiotics cassette of bYS763 was looped out using a cre-expressing plasmid pDR244.

**bYS959** harboring *ΔprkA::cat* was generated upon transformation of PY79 with a Gibson assembly consisting of three fragments: (1) PCR with primers oYS1182 and oYS1183 and PY79 template genomic DNA (containing the upstream of *prkA* locus); (2) PCR with primers oJM028 and oJM029 and chloramphenicol-resistance cassette *loxP-cat-loxP* template (amplified from pWX465); (3) PCR with primers oYS1184 and oYS1185 and PY79 template genomic DNA.

**bYS963** harboring *ΔprkD::cat* was generated upon transformation of PY79 with a Gibson assembly consisting of three fragments: (1) PCR with primers oYS1188 and oYS1189 and PY79 template genomic DNA (containing the upstream of *prkD* locus); (2) PCR with primers oJM028 and oJM029 and *loxP-cat-loxP* template (pWX465); (3) PCR with primers oYS1190 and oYS1191 and PY79 template genomic DNA.

**bYS967** harboring *ΔyabT::cat* was generated upon transformation of PY79 with a Gibson assembly consisting of three fragments: (1) PCR with primers oYS1194 and oYS1195 and PY79 template genomic DNA (containing the upstream of *yabT* locus); (2) PCR with primers oJM028 and oJM029 and *loxP-cat-loxP* template (pWX465); (3) PCR with primers oYS1196 and oYS1197 and PY79 template genomic DNA.

**bYS1053** harboring *ΔclpC::cat* was generated upon transformation of PY79 with a Gibson assembly consisting of three fragments: (1) PCR with primers oYS1341 and oYS1342 and PY79 template genomic DNA (containing the upstream of *clpC* locus); (2) PCR with primers oJM028 and oJM029 and *loxP-cat-loxP* template (pWX465); (3) PCR with primers oYS1343 and oYS1344 and PY79 template genomic DNA.

**bYS1055** harboring *ΔclpP::cat* was generated upon transformation of PY79 with a Gibson assembly consisting of three fragments: (1) PCR with primers oYS1345 and oYS1346 and PY79 template genomic DNA (containing the upstream of *clpP* locus); (2) PCR with primers oJM028 and oJM029 and *loxP-cat-loxP* template (pWX465); (3) PCR with primers oYS1347 and oYS1348 and PY79 template genomic DNA.

**bYS1057** harboring *ΔyrzL::cat* was generated upon transformation of PY79 with a Gibson assembly consisting of three fragments: (1) PCR with primers oYS1349 and oYS1350 and PY79 template genomic DNA (containing the upstream of *yrzL* locus); (2) PCR with primers oJM028 and oJM029 and *loxP-cat-loxP* template (pWX465); (3) PCR with primers oYS1351 and oYS1352 and PY79 template genomic DNA.

**bYS1059** harboring *ΔypiB::cat* was generated upon transformation of PY79 with a Gibson assembly consisting of three fragments: (1) PCR with primers oYS1353 and oYS1354 and PY79 template genomic DNA (containing the upstream of *ypiB* locus); (2) PCR with primers oJM028 and oJM029 and *loxP-cat-loxP* template (pWX465); (3) PCR with primers oYS1355 and oYS1356 and PY79 template genomic DNA.

**bYS893** harboring *ΔprkC, murAA-mNeonGreen* was generated upon transformation of bYS542 with genomic DNA from bYS395. The *loxP-cat-loxP* antibiotics cassette was subsequently looped out using pDR244 plasmid.

**bYS894** harboring *ΔprpC, murAA-mNeonGreen* was generated upon transformation of bYS543 with genomic DNA from bYS395. The *loxP-cat-loxP* antibiotics cassette was subsequently looped out using pDR244 plasmid.

**bYS1061** harboring *amyE::Pxyl-YrzL(T7A)::erm* was generated upon transformation of PY79 with a Gibson assembly consisting of three fragments: (1) PCR with primers oMD191 and oMD226 and bYS545 template genomic DNA; (2) PCR with primers oYS1357 and oYS1358 and PY79 template genomic DNA; and (3) PCR with primers oMD196 and oMD197 and PY79 template genomic DNA.

**bYS1062** harboring *amyE::Pxyl-YrzL(T7E)::erm* was generated upon transformation of PY79 with a Gibson assembly consisting of three fragments: (1) PCR with primers oMD191 and oMD226 and bYS545 template genomic DNA; (2) PCR with primers oYS1359 and oYS1360 and PY79 template genomic DNA; and (3) PCR with primers oMD196 and oMD197 and PY79 template genomic DNA.

**bYS1063** harboring *murAA-mNeonGreen, AmyE::Pxyl-prkC::kan*, was generated upon transformation of bYS397 with genomic DNA from bYS539.

**bYS1065** harboring *murAA-mNeonGreen AmyE::Pxyl-prpC::erm*, was generated upon transformation of bYS397 with genomic DNA from bYS545.

**bYS1067** harboring *murAA-mNeonGreen, ΔclpC::cat* was generated upon transformation of bYS397 with genomic DNA from bYS1053.

**bYS1069** harboring *murAA-mNeonGreen, ΔyrzL::cat* was generated upon transformation of bYS543 with genomic DNA from bYS1057.

**bYS1071** harboring *murAA-mNeonGreen, ΔypiB::cat* was generated upon transformation of bYS542 with genomic DNA from bYS1059.

**bYS1073** harboring *murAA-mNeonGreen, AmyE::Pxyl-yrzL(T7A)::erm, ΔyrzL::cat* was generated upon transformation of bYS1069 with genomic DNA from bYS1057.

**bYS1074** harboring *murAA-mNeonGreen, AmyE::Pxyl-yrzL(T7E)::erm* was generated upon transformation of bYS1069 with genomic DNA from bYS1063.

**bYS1084** harboring *murAA-mNeonGreen, ΔprpC, AmyE::Pxyl-yrzL(T7A)::erm* was generated upon transformation of bYS894 with genomic DNA from bYS1061.

**bMD514** harboring *sfGFP-murG::cat* was generated upon transformation of PY79 with a four-piece Gibson assembly reaction, that contained the following PCR products. (1) PCR with primers oMD349 and oMD350 and PY79 template genomic DNA; (2) PCR with primers oJM028 and oJM029 and *loxP-cat-loxP* template (amplified from pWX465); (3) PCR with primers oMD351 and oMD352 and sfGFP template genomic DNA; and (4) PCR with primers oMD353 and oMD354 and PY79 template genomic DNA.

